# Genomes of *Aegilops umbellulata* provide new insights into unique structural variations and genetic diversity in the U-genome for wheat improvement

**DOI:** 10.1101/2024.01.13.575480

**Authors:** Jatinder Singh, Santosh Gudi, Peter J. Maughan, Zhaohui Liu, James Kolmer, Meinan Wang, Xianming Chen, Matthew Rouse, Pauline Lasserre-Zuber, Helene Rimbert, Sunish Sehgal, Jason Fiedler, Frédéric Choulet, Maricelis Acevedo, Rajeev Gupta, Upinder Gill

## Abstract

*Aegilops* spp. serve as an important reservoir for novel sources of resistance or tolerance to biotic and abiotic stresses. To harness this reservoir, we have generated a high-quality chromosome-level genome assembly of an *Ae*. *umbellulata* accession PI 554389 using a combination of PacBio HiFi, Oxford nanopore, and chromosome conformation capture (Hi-C) sequencing technologies and resequenced 20 *Ae. umbellulata* genomes using Illumina sequencing. We assembled a 4.20 Gb genome spanned over seven chromosomes, rich in repetitive elements (∼84%), achieving a QV of 59.54 with 98.14% completeness. The phylogenetic analysis places the U-genome with D-lineage, but major and distinct rearrangements were revealed in the U-genome. Unique transposon landscape of diploid U-genome and complex chromosomal rearrangements, most prominently in 4U and 6U chromosomes uncovered a distinct evolutionary trajectory of *Ae. umbellulata*. Additionally, the resequencing of geographically and morphologically diverse *Ae. umbellulata* accessions revealed three distinctive evolutionary sub-populations. Resequencing also identified six new haplotypes for *Lr9*, the first leaf rust resistance gene introgressed and cloned from *Ae. umbellulata.* These genomics resources along with high levels of resistance in the resequenced accessions against five devastating wheat diseases affirmed the genetic potential of *Ae. umbellulata* for wheat improvement.

## Main

Wheat wild relatives (WWRs) serve as an important source of stress resiliency traits for wheat improvement. *Aegilops umbellulata* Zhuk. is one such diploid (2n=2×=14) WWR with a large and repetitive U-genome similar to other *Triticeae* genomes^1,2^. *Ae*. *umbellulata*, first reported by Zhukovsky (1928), is a self-pollinated annual grass species that grows primarily in the subtropical ecosphere. It is distributed more prominently in Turkey but has also spread to other West Asian countries along the Fertile Crescent, including Iraq, Lebanon, Iran (West), Syria (North); and Caucasus region – Azerbaijan and Armenia^3,4^. Despite its narrow geographical distribution, *Ae. umbellulata* harbors substantial genetic diversity which is relatively higher than the *Ae. tauschii*, the D-genome donor of cultivated wheat^5^. *Ae. umbellulata* played a key role as a diploid progenitor in the evolution of seven polyploid species belonging to section *Pleionathera*, such as *Ae. triuncialis* L. (U^t^C^t^), *Ae. peregrina* Hack. (U^p^S^p^), *Ae. kotschyi* Boiss. (U^k^S^k^), *Ae. columnaris* Zhuk. (U^c^X^c^), *Ae. geniculata* Roth. (U^g^M^g^), *Ae. biuncialis* Vis. (U^b^M^b^), *Ae. neglecta* Req. ex Bertol. (U^n^M^n^), and *Ae. neglecta* (U^n^M^n^N^n^)^6^.

Multiple genome analysis studies have been conducted to understand the evolution and dynamics of the U-genome in diploid and polyploid species^6–8^. For instance, Kimber and Yen (1989) explored the chromosomal pairing in interspecies hybrids between autotetraploid, *Ae. umbellulata* (UUUU) and the different natural polyploid species containing the U-genome. Similarly, Badaeva and co-authors (2004) employed heterochromatin banding patterns such as C-banding and fluorescence in situ hybridization (FISH) to study the phylogenetic relationship between *Ae. umbellulata* and the seven polyploid species sharing the U-genome. These studies revealed that the U-genome in all tetraploid and hexaploid species is derived from *Ae. umbellulata* and is highly stable in all polyploid species. However, the other genomes (viz., C, S, M, or N) present in species belonging to *Pleionathera* section tend to be modified due to genome instability^6^. These genomic instabilities might occurred due to the meiotic instability induced during homoeologous chromosome pairing in these polyploids^9^ or due to increased activity of transposable elements (TEs)^10–12^. Badaeva and co-authors (2004) hypothesized that the U-genome from *Ae. umbellulata* might have specific classes of TEs, which are responsible for inducing chromosomal rearrangements in other genomes upon polyploidization, leading to larger modifications in the C, S, M, or N genomes of allopolyploids containing the U-genome.

Due to high genetic diversity for resilience to multiple biotic and abiotic stresses, several efforts have been made to introgress agronomically important traits, including stress resilience and improved baking quality, from *Ae. umbellulata* into cultivated wheat^13–20^. In 1950s, first resistance gene, *Lr9, was* transferred from *Ae. umbellulata* to cultivated wheat through X-ray irradiation^13^. Subsequently, other resistance genes from *Ae. umbellulata*, including *Lr76* (leaf rust), *Yr70* (stripe rust), and *PmY39* (powdery mildew) were successfully introgressed into wheat^18,21^. Resistance to destructive stem rust races TTTTF and TTKSK (Ug99) was recently reported in *Ae. umbellulata* and a major quantitative trait loci (QTL) was mapped on chromosome 2U^15^. Besides disease resistance, allosyndetic pairing was induced between chromosome 1U of *Ae. umbellulata* and group 1 chromosomes of wheat to transfer high-molecular weight glutenin subunit (HMW-GS) genes (*1Ux* and *1Uy*)^19,22^. Song et al. (2023), recently reported that, *Ae. umbellulata* has high zinc, iron, and seed gluten content compared to hexaploid wheat and *Ae. tauschii*. The study also revealed higher levels of resistance to wheat stripe rust^23^. In addition to fungal pathogens, *Ae. umbellulata* was found to carry resistance to Hessian fly^24^. Despite its huge potential to improve wheat, the genomic resources for *Ae. umbellulata* are not well established to effectively harvest novel traits/genes residing in this species except for recently published *Ae. umbellulata* genome in a parallel study^25^.

Here we presented a comprehensive analysis of the structural variations and the evolution of U-genome by generating a high-quality chromosome scale assembly of *Ae. umbellulata* accession PI 554389 that has been shown to resistant to multiple diseases. Comparative genome analysis across the *Triticeae* group revealed major chromosomal rearrangements unique to *Ae. umbellulata* which could have implication in pre-breeding and introgressions efforts. Furthermore, we resequenced 20 geographically and morphologically diverse *Ae. umbellulata* accessions with varying levels of resistance to multiple wheat diseases including, leaf rust, stripe rust, stem rust, tan spot, and bacterial leaf streak (BLS) to understand the genetic diversity in *Ae. umbellulata* species. Additionally genotypic and phenotypic evaluation revealed the presence of six allelic variants of *Lr9* (the only leaf rust resistance gene cloned from *Ae. umbellulata* to date), along with additional leaf rust resistance genes, in these 20 accessions.

## Results

### Telomere-to-telomere genome assembly and gene annotations of *Aegilops umbellulata*

*Aegilops umbellulata* accession, “PI 554389” was used to generate a telomere-to-telomere (T2T) chromosome level genome assembly. This accession was originally collected from Turkey in 1991 and was found to confer high levels of resistance to multiple wheat diseases including leaf rust, stem rust, and tan spot (Fig. 1a). A total of 4.28 Gb of *Ae. umbellulata* genome was assembled using: (i) PacBio HiFi reads (126.7 Gb) generated by circular consensus sequencing (CCS); (ii) Nanopore long reads (12.81 Gb) generated by Oxford Nanopore MinION sequencing; and (iii) High-throughput chromosome conformation capture (Hi-C) derived short-reads (Supplementary Fig. 1a; Supplementary tables 1-3). Initially a primary contig assembly was generated using the hifiasm assembler and HiFi data only. The primary contig assembly consisted of 1389 contigs with an N50 and L50 of 8.79 Mb and 141, respectively. Hi-C scaffolding of the primary contig assembly yielded seven pseudomolecules covering 4.20 Gb with N50 value of 634.10 Mb (Table 1; Supplementary table 4). Additionally, PacBio and ultra-long Nanopore reads were used to close 252 gaps present in seven pseudomolecules (Supplementary table 5). The pseudomolecules were oriented and assigned chromosomes names based on their homology with the IWGSC RefSeq_v2.1 D sub-genome^26^ (Fig. 1b; Supplementary table 6). Of the total seven chromosomes, we found telomeres on both ends for four chromosomes (4U, 5U, 6U, and 7U) and one end for remaining three chromosomes (1U, 2U, and 3U) (Supplementary Fig. 1b). Chromosome sizes ranged from 488.85 Mb (1U) to 658.84 Mb (6U) and the GC content ranged from 46.91% (7U) to 47.3% (6U) (Fig. 1b; Supplementary table 7; 8). The complete chloroplast and mitochondrial genomes were also de novo assembled (Supplementary Fig. 1c) and any unanchored contigs that aligned to these extranuclear genomes were removed from the final assembly. Quality assessment of the assembled nuclear genome using INSPECTOR revealed a 99.99% mapping rate of HiFi reads with genome quality score of 59.54. Genome completeness was assessed using BUSCO, which identified that >98% of single copy conserved orthologs within the embryophyta dataset were identified as complete, with less than 5% being duplicated, as expected for a diploid species (Supplementary Fig. 1d; Supplementary table 9).

**Figure 1.**
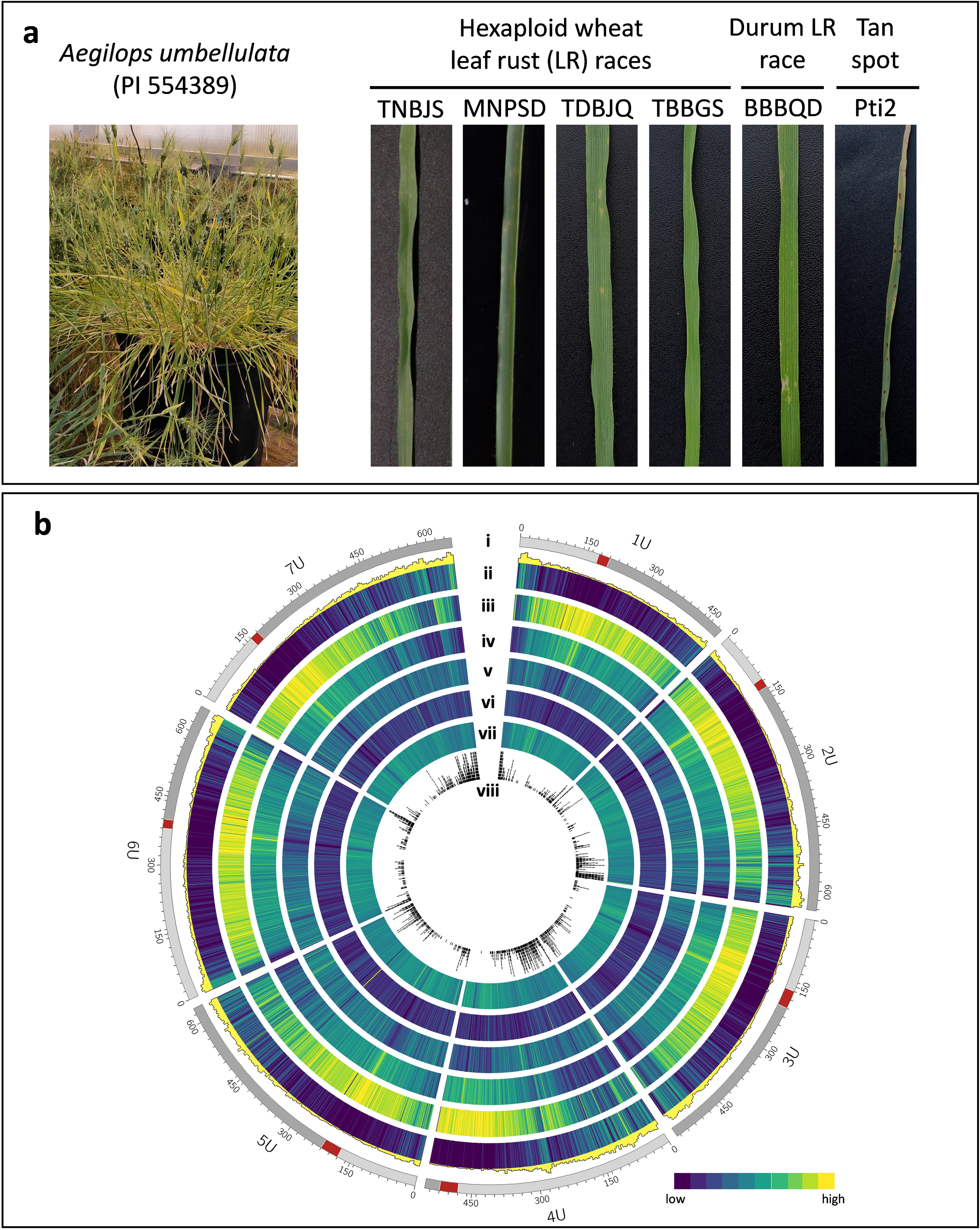
Genome sequencing of *Aegilops umbellulata* accession, PI 554389**: (a)** phenotypic appearance and the disease reaction of *Ae. umbellulata* accession, PI 554389 to leaf rust and tan spot disease. TNBJS, MNPSD, TDBJQ, and TBBGS are the leaf rust races affecting *Triticum aestivum*, BBBQD is the leaf rust races affecting durum wheat, and Pti2 is the race causing tan spot in wheat; **(b)** Circos plot showing the genomic features of *Ae. umbellulata* genome. Within the circos plot: **circle-i** depicts the individual *Ae. umbellulata* chromosomes with their physical positions (Mb); **circle-ii** depicts the number (histogram) and density (heatmap) of high-confidence genes; **circle-iii** depicts the percentage distribution of transposable elements (TEs); **circle-iv** depicts the percentage distribution of Gypsy (RLG) elements; **circle-v** depicts the percentage distribution of Copia (RLC) elements; **circle-vi** depicts the percentage distribution of CACTA (DTC) elements; **circle-vii** depicts the percentage of GC content; and **circle-viii** depicts the number of resistant gene analogs (RGA).

**Table 1:**
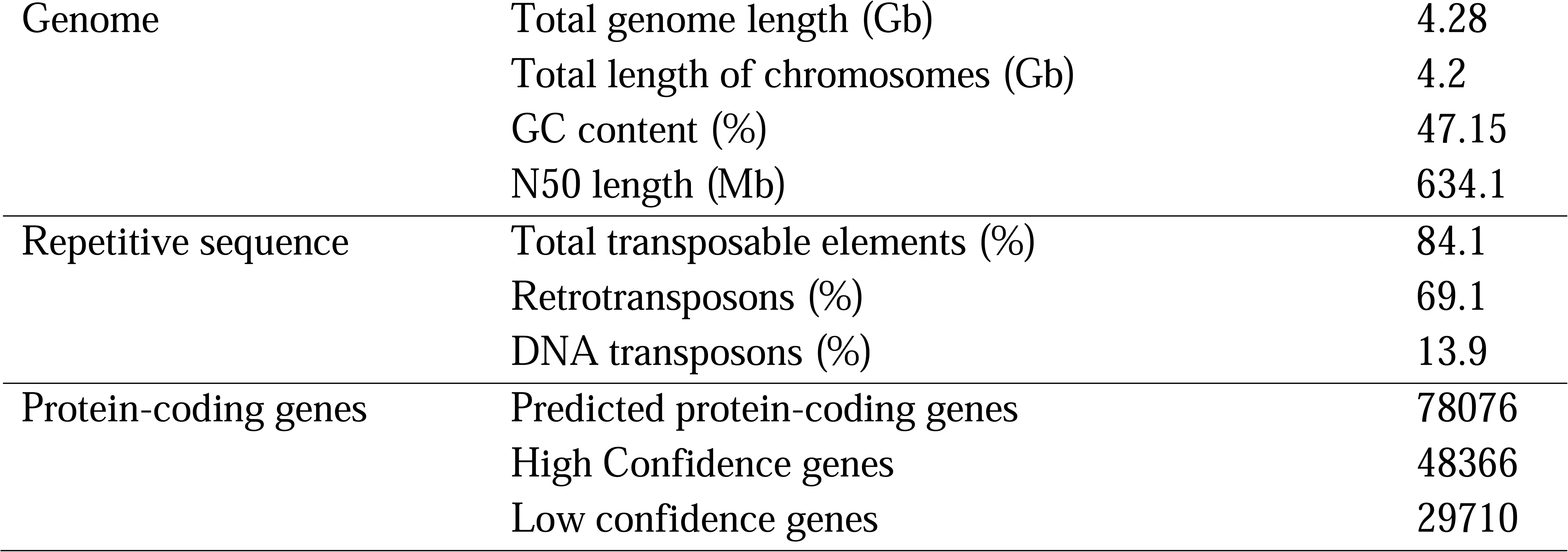
Statistics of genome features of *Ae. umbellulata* chromosome level assembly.

The full-length transcript data was generated using PacBio iso-seq single molecule real-time (SMRT) sequencing technology by pooling RNA samples from four different tissues (seedling shoots, seedling roots, leaves after vernalizations, and immature spikes) and used as primary evidence to annotate 78,076 gene models, of which, 48,366 were high-confidence (HC) genes (Table 1; Fig. 1b; Supplementary table 7). Among the HC genes, a total of 2,162 were predicted to be resistance gene analogs (RGAs),including 790 nucleotide-binding domain leucine-rich repeats (NLRs), 1,094 receptor-like kinases (RLKs), 160 transmembrane-coiled-coil (TM-CC), and 117 receptor-like proteins (RLPs)^27^ (Supplementary table 10).

### Unique transposon landscape shapes the evolutionary trajectory of *Ae. umbellulata* species

The transposable elements (TEs) search identified 1.24 million copies of TEs in the U-genome accounting for 3.6 Gb. This proportion (84.1% of U-genome) is very similar to previous estimates in A-B-D sub-genomes of *T. aestivum* and other related tetraploid and diploid *Triticeae* species, where 83 to 88% repetitive sequence are reported (Supplementary Fig. 2)^28^. At the superfamily level, the TE types are similar to what were previously observed for A-B-D sub-genomes of wheat. Gypsy, Copia and CACTAs represent 47.1%, 17.7% and 12.7% of total TEs, respectively, while the remaining superfamilies are much less abundant (Supplementary Table 11). Among the 291 TE families (representing at least 100 Kb) shaping the U-genome, only 15 (5%) exhibit a substantial change in abundance (log2 fold-change <-2 or >2) between U and at least one of the compared genomes: five LTR-retrotransposons (RLG_famc25-27-36, RLC_famc26, and RLX_famc21), one LINE (RIX_famc40), two DNA transposons (DTM_famc22 and DXX_famc3), and seven unclassified repeats (XXX_famc10-24-46-47-53-57-81). These 15 families are low copy number with abundance close to the 100 Kb cutoff. The MITE Mutator DTM_famc22 (named Gabriel in TREP^29^) is overrepresented in the U-genome because of the presence of a 2 Mb cluster on 2U consisting of 939 tandemly repeated copies of this ∼400 bp MITE. Except these particular cases, the data broadly suggests that 95% of the families are present in proportions that are stable since U-A-B-D lineages diverged, although they have experienced a near-complete TE turnover. Such dynamics confirms previous conclusions achieved by comparing distant^30^ and closely related genomes to trace novel insertions^28^. The TE turnover was not accompanied by lineage-specific TE burst/loss. In general, TE families tend to maintain at a relatively constant copy number and the U-genome is another example of *Triticeae* that followed this equilibrium model of evolution.

### Comparative genome analysis revealed major chromosomal rearrangements unique to U-genome

Comparative genome analysis is a powerful tool to dissect the genomic architecture and the evolutionary history of species. Rooted species tree analysis using multi-copy orthologous genes among 11 *Triticeae* species and one outgroup species revealed three clades belonging to A-, B-, and D-lineages, where *Ae. umbellulata* was phylogenetically placed in D-lineage indicating a shared ancestry (Fig. 2a; Supplementary Table 12). Among the D-lineage species, *Ae. umbellualata* (U-genome) diverged earlier followed by the separation of the D-and S-genome species from a common ancestral parent. During this evolutionary separation, the U-genome accumulated major chromosomal rearrangements^31^ which led to broader genetic diversity in the U-genome^5^, as evident from the morphological and genetic variations in *Ae. umbellulata*. To further understand these structural variances, we performed a whole genome comparative analysis of the U-genome within the D lineage species and outside of the D lineage, including *Ae. speltoides*, *T. urartu*, A-B-D sub-genomes of wheat and phylogenetically distant species *H. vulgare* and *B. distachyon*. We observed high levels of chromosomal discordance amongst studied species relative to the U-genome (Fig. 2b; Supplementary Fig. 3). To study these structural variations and evolutionary relationship of the U-genome with A-, B-, and D-lineages, we performed the macro-synteny analysis of *Ae. umbellulata* with *T. urartu*, *Ae. spletoides*, *Ae. tauschii*, *Ae. sharonensis*, and *Ae. longissima*. Our results revealed major chromosomal rearrangements unique to the U-genome including several prominent non-reciprocal translocations in 4U and 6U chromosomes (Fig. 2c; Supplementary Fig. 4). Based on our synteny analysis among *Ae. tauschii*, *Ae. umbellulata,* and *Ae. longissima*, we identified a unique 7S^l^/4S^l^ translocation (Fig. 2c). This translocation has been reported previously only in *Ae. longissima* but absent in the rest of the S and D genome species^2^. We also observed 5U/6U reciprocal translocation in *Ae. umbellulata* compared to other species except for *T. urartu* (Supplementary Fig. 4). In addition to translocations, two major intra-chromosomal inversions on 7U and 2U chromosomes and multiple inter-chromosomal translocated inversions were detected (Fig. 2c). In addition to intra-chromosomal inversion in 7U, a non-reciprocal translocation from group 3 chromosome was also observed. In contrast, chromosomes 1U, 3U and 5U remained remarkably conserved across the U-, A-, B-, D-, and S-genomes. All these structural variations lead to the formation of a unique U-genome of the *Ae. umbellulata* with higher genetic diversity compared to other *Triticeae* species^5,32^.

**Figure 2.**
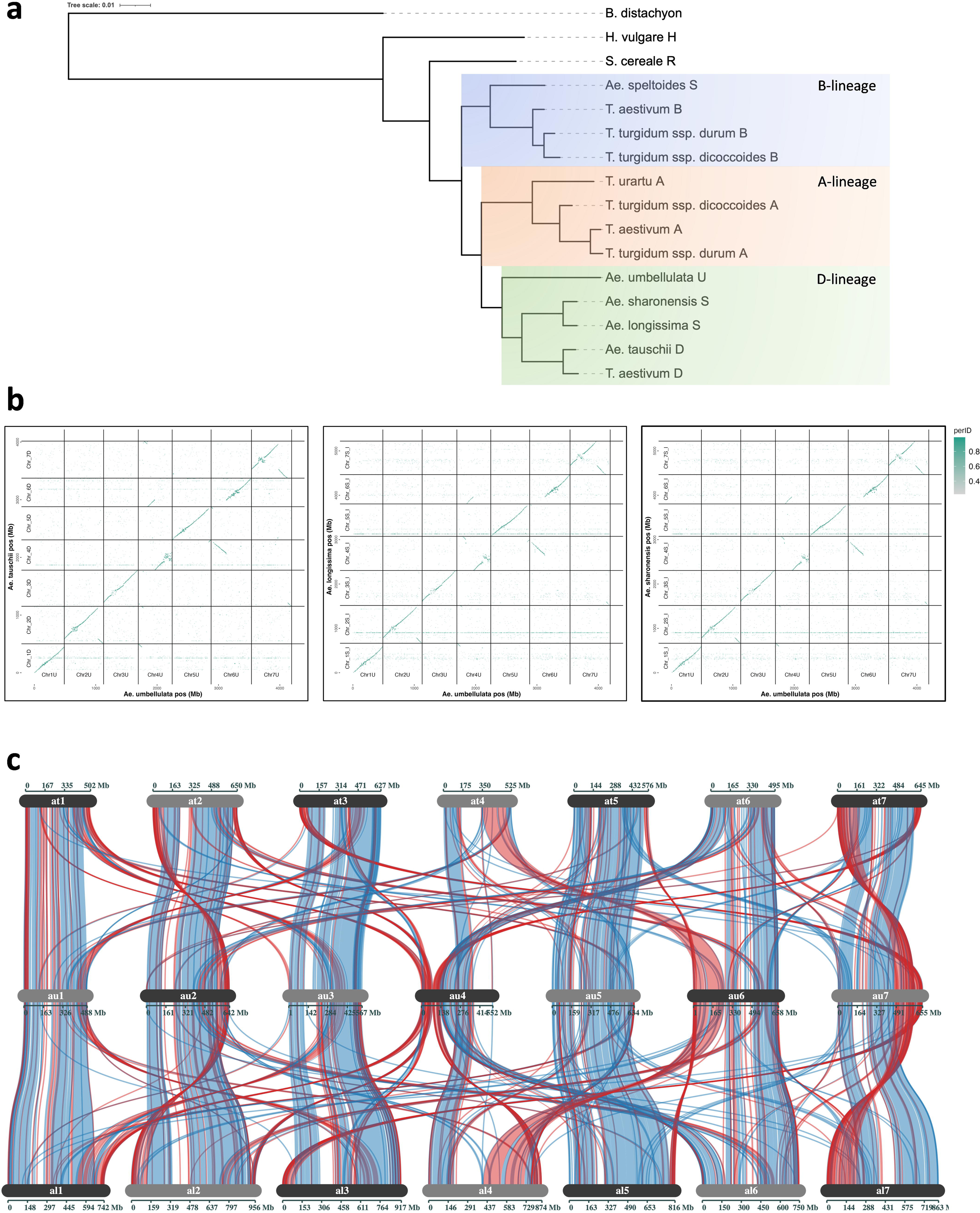
Comparative genome analysis of *Aegilops umbellulata* with other species: **(a)** dendrogram showing the phylogenetic relationship of *Ae. umbellulata* with 10 *Triticae* spp. and one outgroup species *Brachypodium distachyon*; **(b)** whole genome comparison of *Ae. umbellulata* with *Ae. tauschii*, *Ae. longissima*, and *Ae. sharonensis*; **(c)** synteny and collinearity analysis of *Ae. umbellulata* with *Ae. tauschii* and *Ae. longissima*.

### Complex chromosomal rearrangements drive the evolution of present day 4U and 6U chromosomes

We observed pronounced chromosomal rearrangements particularly in chromosomes 4U and 6U. Therefore, chromosomes 4U and 6U were examined in detail for their collinearity with two D lineage species, *Ae. tauschii* (D) and *Ae. longissima* (S^l^) (Fig. 3a; Supplementary Fig. 5). The previously known 7S^l^/4S^l^ translocation in *Ae. longissima*, prompted us to study the S^l^ genome along with the D genome of *Ae. tauschii*^2,33,34^. Our results suggest the presence of ancestral chromosomal segments of 1L (long arm), 2L, 7L, and 6S (short arm) on 4U and segments of 5L and 4L on 6U (Fig. 3a; Supplementary Fig. 5). Based on these findings, we proposed a model to explain the evolution of the 4U and 6U chromosomes (Fig. 3b,c). In the first translocation event, a significant portion of ancestral chromosome 6S (∼120 Mb) may have fused to chromosome 4S. In the subsequent non-reciprocal, inter-chromosomal translocation events, distal ends of ancestral chromosomes 7L (∼55 Mb), 2L (∼34 Mb), and 1L (∼55 Mb) inversely fused to chromosome 4 of *Ae. umbellulata*, which gave birth to the modern day 4U chromosome of *Ae. umbellulata* (Fig. 3b). Similarly, 6U chromosome may have evolved through non-reciprocal translocation of a substantial region of the ancestral 4L chromosome (∼210 Mb) into broken/wounded chromosome 6S end resulted from translocation of 6S segment to 4S (Fig. 3c). These events suggest that 4U and 6U have evolved via independent sequential translocation events rather than a single major translocation event. Additionally, the 4L segment incorporated into the 6U chromosome underwent a reciprocal translocation with the 5U chromosome, exchanging a region of ∼35 Mb. Moreover, a paracentric inversion in a very small portion (∼2-3 Mb) of chromosome 4 segment present in 6U occurred. In addition to these translocations and inversions, there were multiple intra-chromosomal rearrangements during the evolution of 4U and 6U chromosomes.

**Figure 3.**
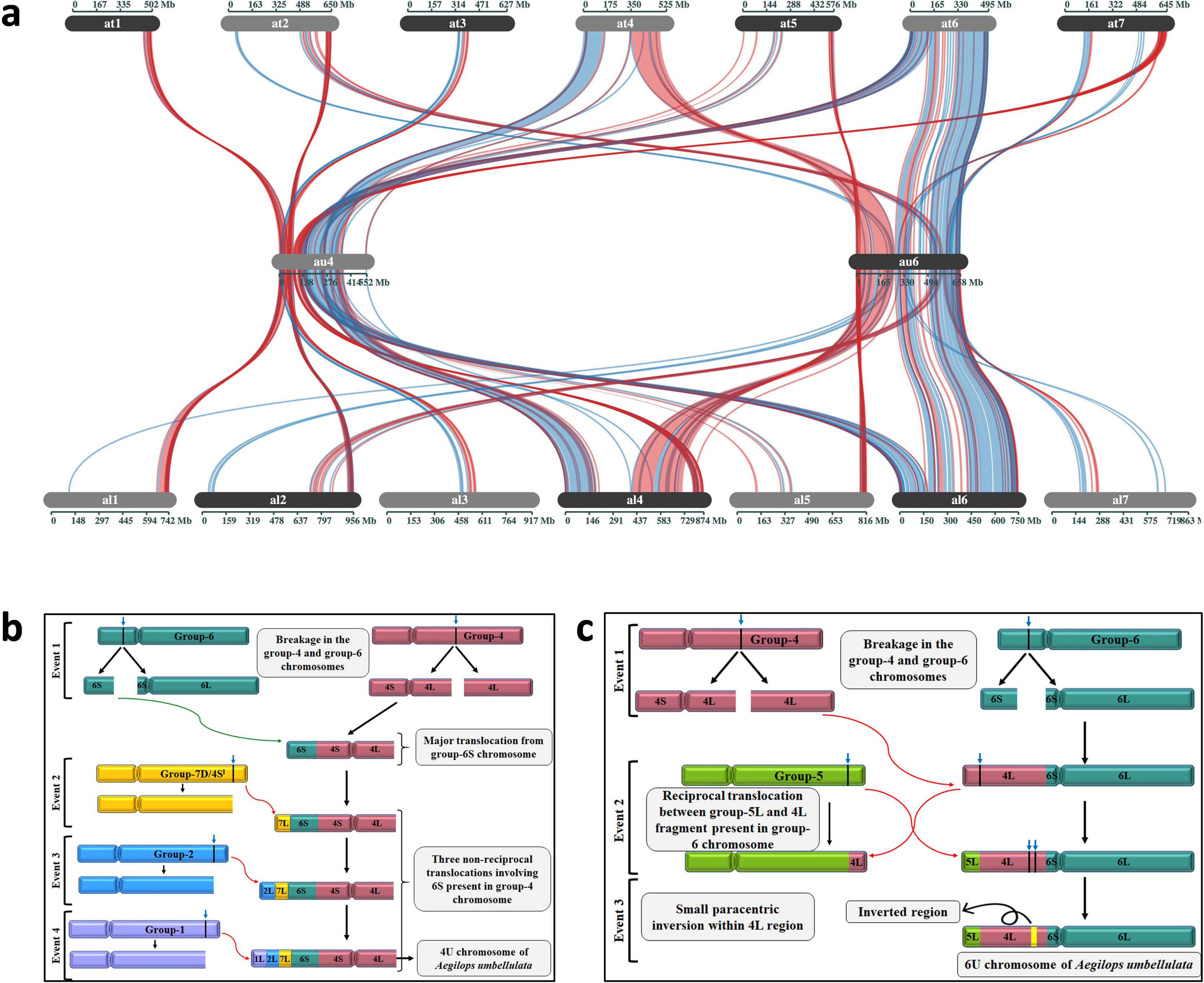
Evolution of 4U and 6U chromosomes of *Aegilops umbellulata*: **(a)** synteny and collinearity analysis of 4U and 6U chromosomes of *Ae. umbellulata* (au) with all chromosomes of *Ae. tauschii* (at) and *Ae. longissima* (al); **(b)** model depicting the origin of the 4U chromosome of *Ae. umbellulata*; **(c)** model depicting the origin of the 6U chromosome of *Ae. umbellulata*.

### Resequencing of *Ae. umbellulata* reveals rich genetic diversity in U-genome

To investigate the genetic diversity in *Ae. umbellulata*, the whole genomes of 20 accessions were resequenced at ∼10X coverage. The selected accessions display high variability for morphology and resistance to five wheat diseases, such as leaf rust, stripe rust, stem rust, tan spot, and BLS (Fig. 4a, Supplementary Table 13, Supplementary Fig. 6-9) and represent a large geographical distribution (Fig. 4b). Variant calling using GATK (v.4.1.8.0) identified a total 86,931,487 SNPs among the 20 accessions (Supplementary table 14). After hard masking and filtering SNPs for SNP depth (4 < DP >15), SNP cluster (>3 SNPs within 10lJbp window), non-biallelic SNPs, and SNPs unanchored to chromosomes with missing value of 90%, a total of 7,184,562 SNPs were retained and used for further analyses (Supplementary table 14). Principal component analysis (PCA) using the filtered sets of SNPs divided 20 accessions into three sub-groups (Fig. 4c). Fifteen accessions were present in group-I, whereas only three and two accessions were present in the group-II and group-III, respectively. Most of the accessions collected from Turkey and all accessions from other regions fell in the group-I, except for the accessions collected from Serbia (group-II) and the United Kingdom (group-III). Accessions in the group-I have comparatively higher level of resistance to various leaf rust races, whereas the accessions in group-II and III were susceptible to several leaf rust races (Fig. 4a; Supplementary Table 13). PCA results were further confirmed by estimating the maximum likelihood method for genotype grouping and ancestry coefficients using cross-entropy values, which revealed the presence of three sub-groups (K = 3) (Fig. 4d-f). Nucleotide diversity (*π*) analysis among 20 accessions as well as within each sub-group revealed the maximum genetic diversity in group-III (*π* = 0.00035) and minimum genetic diversity in group-I (*π* = 0.0001) (Fig. 3g). Additionally, the pairwise nucleotide diversity (*π*) analysis revealed certain regions within each chromosome with very low and high nucleotide diversity (Supplementary Fig. 10).

**Figure 4.**
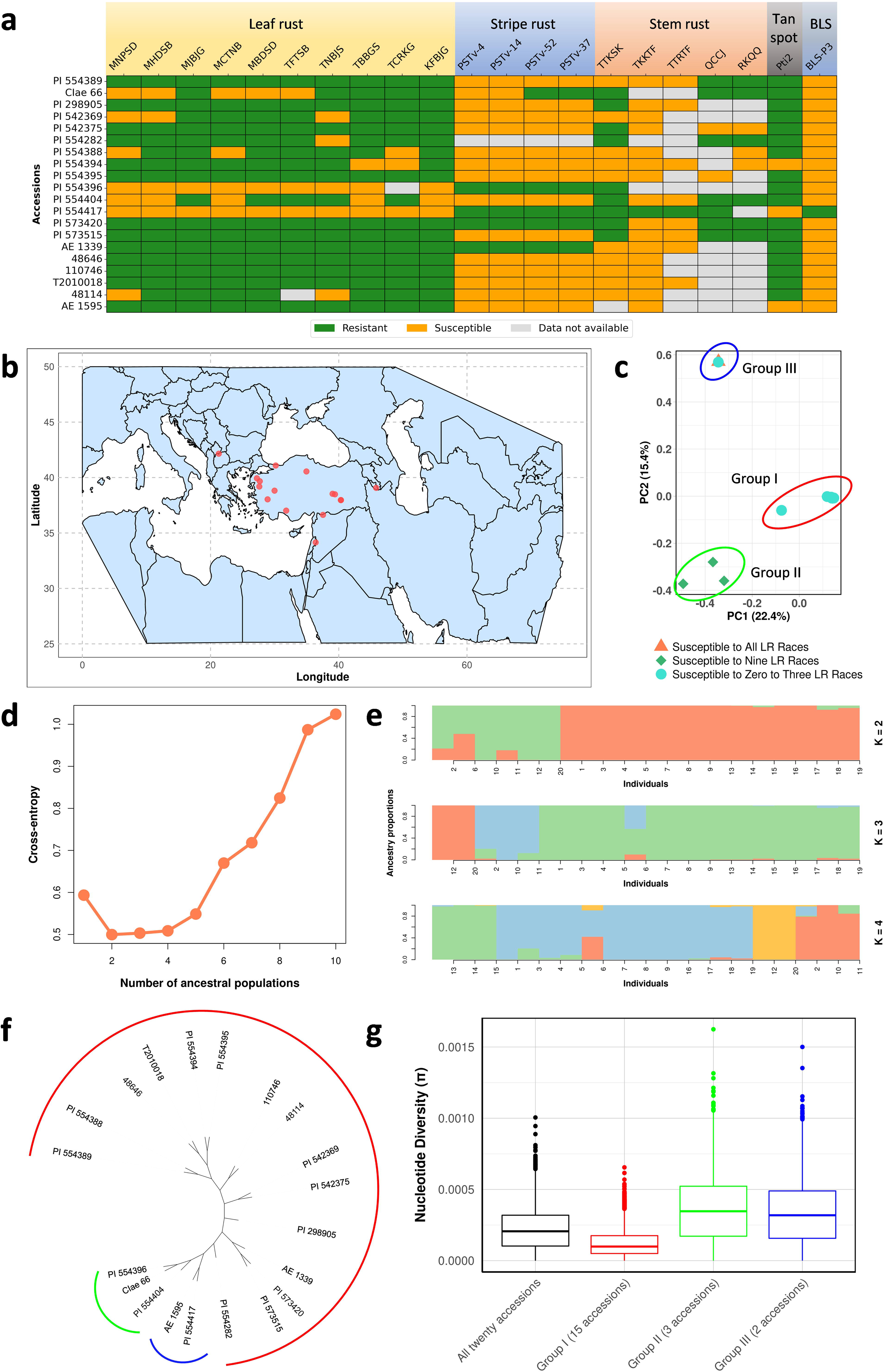
Assessing genetic diversity among 20 *Aegilops umbellulata* accessions: **(a)** disease reaction of 20 *Ae. umbellulata* accessions for multiple races of leaf rust (10 races), stripe rust (4 races), stem rust (5 races), tan spot (1 race), and bacterial leaf streak (BLS); **(b)** geographical distribution of 20 *Ae. umbellulata* accessions; **(c)** principal component analysis (PCA) to study the population stratification among *Ae. umbellulata* accessions. PCA divided all accessions into three subgroups with 15, 3 and 2 accessions in each subgroup; **(d)** cross entropy values showing the number of ancestral populations among the *Ae. umbellulata* accessions; **(e)** landscape and ecological association (LEA) analysis for studying the population diversity among *Ae. umbellulata* accessions. Where, K is the number of subgroups in the population; **(f)** maximum-likelihood method for grouping *Ae. umbellulata* accessions into subgroups; **(g)** nucleotide diversity (π) analysis to estimate the amount of genetic diversity residing among all *Ae. umbellulata* accessions as well as within each subgroup.

### Allelic variants of historically important *Lr9* along with novel leaf rust resistance genes

Phenotypic and genotypic evaluations were used to study the presence of *Lr9* and additional leaf rust resistance genes in 20 *Ae. umbellulata* accessions. PI 554389 and seven additional accessions were highly resistant to 10 leaf rust races including the two races (MNPSD and TNBJS) that are virulent on *Lr9* (Supplementary table 13), suggesting the presence of *Lr9* and/or new leaf rust resistance gene(s). Two accessions, PI 554282 and PI 542375 are resistant to all tested leaf rust races except TNBJS and TBBGS, respectively, suggesting the presence of two or more novel *Lr* genes. Only one accession, 48114, has a reaction pattern similar to *Lr9*, while the remaining accessions displayed five unique reaction patterns suggesting the presence of *Lr9* and additional *Lr* genes in *Ae. umbellulata* that are yet to be identified.

Since the *Lr9* gene has been cloned^35^, we searched *Lr9* alleles in the 20 resequenced accessions. A complete copy of *Lr9* was present in 10 accessions including PI 554389 (Supplementary table 15). Based on the amino acid sequence variations from the cloned *Lr9* gene, we categorized the *Lr9* alleles into six haplotypes (Fig. 5a; Supplementary table 15). Remaining accessions, which lack the complete *Lr9* coverage, were also missing the transcripts of *Lr9*, further verifying the absence of *Lr9* (unpublished data). The haplotype analysis of *Lr9* revealed the high conserveness in protein kinase 1 (PK1) domain in all six haplotypes. In contrast, several amino acid substitutions were detected in protein kinase 2 (PK2), Von Willebrand factor type A (vWA), and vWaint domains (Fig. 5a; Supplementary tables 15, 16; Supplementary Fig. 11). A single amino acid substitution from Leucine to Proline (L642P) in the PK2 domain was found in all six haplotypes compared to the cloned *Lr9* (Fig. 5a; Supplementary table 16; Supplementary Fig. 11). It is possible that a natural mutation had occurred in the original source (TA1851) of *Lr9* before it was introgressed into the hexaploid wheat cultivar ‘Thatcher’. The L642P substitution also defines haplotype-I (Hap-I) and is present in three accessions, 48114, 48646, and PI 554395. The phenotypic virulence pattern of 10 leaf rust races also supports the presence of a functional *Lr9* allele in 48114 due to its susceptibility only to *Lr9* virulent races, MNPSD and TNBJS (Supplementary table 13). Therefore, despite the single amino acid substitution, Hap-I is a functional allele of *Lr9*. In contrast, the other two accessions, 48646 and PI 554395, harboring Hap-I are resistant to all tested leaf rust races, including the *Lr9* virulent races, MNPSD and TNBJS, suggesting the presence of a novel leaf rust gene(s) along with *Lr9* in these accessions. Hap-II, represented by PI 542369, has a single amino acid insertion of threonine (S21_S22insT) along with the aforementioned L642P substitution (Fig. 5a; Supplementary table 16; Supplementary Fig. 11). PI542369 displayed a leaf rust reaction similar to 48114 except an additional susceptibility to MHDSB which suggest that Hap-II may also be a functional allele of *Lr9*. Hap-III, Hap-IV, Hap-V, and Hap-VI have amino acid substitutions mainly in PK-2 or PK-2 and vWA or PK2, vWA and vWaint domains. Accessions harboring the Hap-III (PI 554389), Hap-IV (110746), Hap-V (PI 298905), and Hap-VI (PI 573515) were completely resistant to all the tested leaf rust races including the races (MNPSD and TNBJS) virulent on *Lr9*. It is possible that these lines have functional alleles of *Lr9* along with an additional *Lr* gene(s) or the unknown *Lr* gene(s) is effective against all tested races. We also identified two accessions (PI 554417 and AE 1595) which had the complete *Lr9* gene, with premature stop codon before PK-1 domain (Supplementary table 15). Both accessions lacked the *Lr9* transcript (unpublished data). PI 554417 was highly susceptible to all tested races and AE 1595 was moderately susceptible to four out of 10 races suggesting the absence of a functional *Lr9* allele in both accessions (Supplementary tables 13, 15).

**Figure 5.**
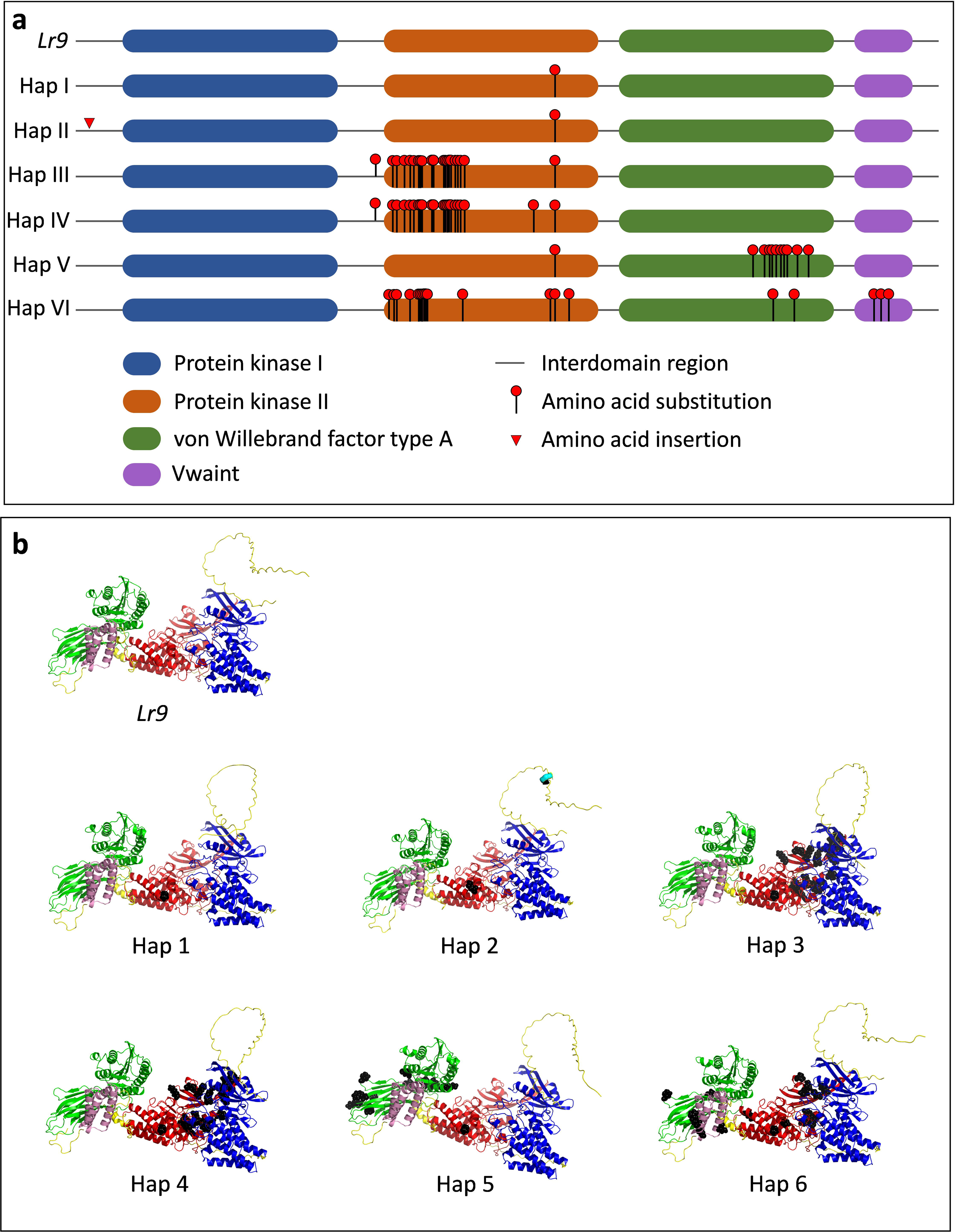
Identifying *Lr9* haplotypes (Hap) from 20 *Aegilops umbellulata* accessions: **(a)** comparing protein sequence of *Lr9* with the allelic variants identified from 20 *Ae. umbellulata* accessions; **(b)** alpha-fold prediction for *Lr9* haplotypes.

## Discussion

The genus, *Aegilops* has been extensively explored to harness useful traits for bread wheat improvements^36^. *Ae. umbellulata* is one of the WWRs that carries large diversity for stress resiliency. At the chromosomal level, *Ae. umbellulata* has structural rearrangements that are unique to this species^31,37^. To study these changes and the associated genetic diversity, a T2T genome assembly along with single-base resolution genomic variation map of 20 resequenced *Ae. umbellulata* accessions was developed in this study. Higher proportion of repetitive elements in *Triticeae* are major drivers that have shaped the genomic landscape of wheat and WWRs^38,39^. In the present study, we found a unique TEs landscape in *Ae. umbellulata*, which has little conserved orthology with wheat TEs. This difference in the distribution of *Ae. umbellulata* TEs is likely to have led to gross chromosomal structural rearrangements that characterize the U-genome. TEs such as RLG_Sabrina, RLG_WHAM, and RLC_Angela are already known to drive species evolution in rye and wheat^39,40^. Phylogenetically, U-genome is grouped in D-lineage, but formed a distinct clade from D– and S-genomes (Fig. 2). A previously known 7S^l^/4S^l^ translocation in *Ae. longissima*^34^ was also observed in *Ae. umbellulata* although in what appears to be a distinct and independent event. The rearrangements in chromosome 4U and 6U are especially significant, where major segments of 4U and 6U chromosomes have experienced extensive reciprocal and non-reciprocal intrachromosomal translocation events (Fig. 3). Based on the collinearity analysis with *Ae. tauschii* and *Ae. longissima*, the present day 4U and 6U chromosomes appear massively restructured, consisting of large portions of several chromosomes, specifically 1L-2L-7L-6S-4S-centromere-4L and 5L-4L-6S-centromere-6L, respectively (Fig. 3). The centromeric regions of chromosome 4U and 6U are syntenic to wheat group-4 and 6 chromosomes, respectively. In the absence of a high-quality genome, previous studies have been unable to correctly identify the origin of these chromosomes, resulting in erroneously interchanging their names^25,41^. In the present study we rectified this error and assigned the correct names for these chromosomes to facilitate future genomics studies in *Triticeae*, also suggested by some previous studies^31,42^.

The complex chromosomal rearrangements observed in *Ae. umbellulata* is likely to have played a role in the expanded genetic diversity observed in the species even though it is distributed across a narrow geographical area^5,6,43^. Higher genetic diversity reported in *Ae. umbellulata* compared to *Ae. tauschii* is reflected by significantly higher levels of observed genetic polymorphism as evidenced by increased numbers of identified SNP and the occurrence of alleles with rare frequencies^5,32^. Not unexpectedly, this increase in genetic diversity has also increased the variation observed in plant morphological features such as spike length, spike shape, and plant growth etc. and multifarious reactions for different wheat diseases, *viz.* tan spot, BLS, and three wheat rusts (Fig. 4a; Supplementary Figs. 6-9). These traits can be harnessed to widen the genetic base of modern cultivated bread wheat. Attempts have been made to identify and introgress economically important genes including *Lr9* from *Ae. umbellulata*. Our phenotypic data suggested the presence of potential five to six novel leaf rust genes resistant to prevalent races in the U.S. Furthermore, we identified *Ae. umbellulata* accessions showing resistance to highly virulent global Ug99 and non Ug99 stem rust races (TTKSK, TTRTF, and TKKTF) belonging to different *Pgt* clades I, III, and IV^44–47^. Another widely distributed wheat disease BLS, threatens the wheat production in the US and other parts of the world^48^. Some level of genetic resistance for this disease is found in triticale, but most of the tested wheat germplasm lack resistance for this disease^49^. We identified one highly resistant *Ae. umbellulata* accession, PI 554417, which can be used for future breeding programs to deploy BLS resistance in the cultivated wheat.

In a recent study, Wang et al. (2023) reported the cloning and characterization of a leaf rust resistance gene, *Lr9*. In our study, we identified six haplotypes of *Lr9* located on 4U chromosome. This gene was originally introgressed to wheat on chromosome 6BL from *Ae. umbellulata* by Sears in the 1950s and was reported to be located on chromosome 6U^13,35^. The resulting introgression on chromosome 6BL and aforementioned erroneous naming of chromosome 4U and 6U, might led to the confusion about chromosomal position of *Lr9*. Based on GISH and C-banding patterns, Friebe et al. (1995), reported the transfer of *Lr9* segment in 6B, 4B, 2D, and 7B wheat chromosomes^50^. In the present study we reported that the present day 4U chromosome contains the chromosomal segments from these four chromosomes. We speculate that the introgression of *Lr9* is a homoeologous exchange of chromosomal segments instead of random incorporation^31^. We located *Lr9* at chromosome 4U at 72439902-72455974 bp, which is homoeologous to the long arm of the wheat group-2 chromosome. Moreover, we also reported amino acids substitutions majorly in PK-2 domain of six haplotypes of *Lr9*, which is assumed to be a pseudo-kinase^35^. Our phenotypic and genotypic data supports the presence of at least two functional *Lr9* haplotypes in *Ae. umbellulata* accessions evaluated in this study.

In summary, our study has unveiled major evolutionary changes in U-genome over millions of years, which have resulted in enriched genetic diversity in this species. The genomic resources developed in this study provides a portal to harness the novel traits to build future proof wheat cultivars with elevated resilience to biotic and abiotic stresses, which are posing a threat to wheat productivity. The profound chromosomal rearrangements in *Ae. umbellulata* discovered in his study opens new avenues to study the evolution of wheat tertiary gene pool species including the resistance gene evolution in more detail.

## Methods

### Plant material and disease reaction

*Ae. umbellulata* accession, “PI 554389” was selected for whole genome sequencing based on the resistance response to ten hexaploid leaf rust races, one durum leaf rust race, two stem rust races, and one tan spot race (Fig. 1a, Supplementary table 13). The accession was procured from the United States Department of Agriculture-Germplasm Resources Information Network (USDA-GRIN), which was originally collected near the Hazar Lake, 35 km South East of Elazığ, Turkey.

### Genome assembly

#### PacBio HiFi library preparation and DNA sequencing

*Ae. umbellulata* seedling was grown in a controlled growth chamber adjusted with 12 hours (h) photoperiod and day/night temperature set to 20°/18°C. The hydroponic growth solution was made using MaxiBloom® Hydroponics Plant Food (General Hydroponics, Sevastopol, CA, United States) at a concentration of 1.7 g/L. In preparation for PacBio HiFi sequencing, HMW DNA was extracted from two-week-old young leaf tissues given 72-h dark-treatment using a CTAB-Qiagen Genomic-tip protocol as described previously^51^. DNA quantity and quality was checked using Qubit dsDNA HS assay and Nanodrop spectrophotometer, respectively. HMW genomic DNA was sheared to 17Kb on a Diagenode Megaruptor and then made into SMRTbell adapted libraries using SMRTbell Express Template Prep Kit 2.0. Size selection was performed using a Sage BluePippin to select fragments greater than 10kb and then sequenced at the BYU DNA Sequencing Center (Provo, UT, USA) using Sequel II Sequencing Kit 2.0 with Sequencing Primer v5 and Sequel Binding kit 2.2 for 30 h with adaptive loading using PacBio SMRT Link recommendations.

Further, Oxford Nanopore sequencing was used to generate super long reads on MinION sequencer using three FLO-MIN112 flow cells. HMW DNA was used for library preparation using SQK-LSK112 ligation kit for nanopore sequencing. The flow cell was primed according to the manufacturer’s instructions and the library was loaded on the flow cell in a dropwise manner (Supplementary table 2). The raw data was basecalled using Guppy (v6.0.1; Oxford Nanopore Technologies, U.K.).

#### Genome assembly and scaffolding

A primary contig assembly was constructed using the PacBio HiFi CCS reads using Hifiasm v16.1^52^ with default parameters for an inbred species (-l 0). Hi-C was used to scaffold the primary contigs assembly into pseudo-molecules. Hi-C reads were aligned to the primary contig assembly using the Burrows-Wheeler Aligner (BWA^53^). Only paired end reads that were uniquely aligned to contigs were retained for downstream analyses. Contigs were clustered, ordered, and oriented using Proximo^TM^, an adapted proximity-guided assembly platform based on the LACHESIS method with proprietary parameters developed at Phase Genomics^54–56^. Gaps between scaffolds within the scaffolded assembly were filled with 100 Ns.

#### Manual gap closing and chromosome assignment

We used long reads generated by PacBio HiFi sequencing and Nanopore MinION sequencing to close the gaps in pseudomolecules, after adapter trimming and filtering the low-quality reads. Gap closing was performed using command line BLASTN. The PacBio HiFi and long nanopore reads that spanned the whole gap were extracted based on their similarity to the 2kb region on both sides of a gap. These extracted reads spanning the gap were visualized using Geneious Prime 2023.0.4 (https://www.geneious.com) for the confirmation. Then, these sequences were used to fill the gaps manually in the assembly. Custom script with more detail on gap closing is provided in the code availability statement. The gap closed pseudomolecules were assigned with chromosome names based on their synteny to *Triticum aestivum* sub-genome D (IWGSC RefSeq_v2.1) using minimap2 (Li 2018).

### Annotations of gene models and transposable elements

#### Iso-seq and transcriptome assembly

For transcriptome assembly, tissue samples were collected at different growth stages from seedlings, roots, leaf tissues after vernalization, and immature spike in liquid nitrogen. The frozen samples were grounded using mortar and pestle. About 100 µg frozen and grounded tissue was used to extract RNA using Qiagen RNeasy Plant Mini Kit (74904) following the manufacturer’s instructions. The quantity and quality of extracted RNA first tested using Nanodrop spectrophotometer, which was further evaluated with Bioanalyzer. After quality check, RNA from all the samples was pooled in equal molar ratios to synthesize full length complementary DNA (cDNA) using a NEBNext® single cell/low input cDNA synthesis and amplification kit (E6421L) which uses a template switching method to generate full length cDNAs (New England BioLabs, Ipswich, MA, USA). IsoSeq libraries were prepared from the cDNA according to standard protocols using the SMRTbell v3.0 library prep kit (Menlo Park, CA, USA) and sequenced on a single SMRT cell 8M using a PacBio Sequel II the DNA sequencing center at Brigham Young University (Provo, Utah, USA).

#### Genome annotation

Species specific repeats identified by RepeatModeler v1.0.11 and RepeatMasker v4.0.9^57^ was used to identify, classify and mask repeat elements within the assembled genome relative to the Repbase-derived RepeatMasker libraries (Dfam 3.0; 20190227; http://www.girinst.org/). The MAKER3 pipeline^58^ was used to annotate the final genome assembly using as primary evidence the *A. umbellulata* transcriptome (see above) and as alternative evidence the predicted gene models and their translated protein sequences from barley (Hv_Morex.pgsb.Jul2020; https://wheat.pw.usda.gov/GG3/content/morex-v3-files-2021), wheat (https://phytozome-next.jgi.doe.gov/info/Taestivumcv_ChineseSpring_v2_1), and panBrachypodium (https://phytozome-next.jgi.doe.gov/info/BdistachyonPangenome_v1) as well as all proteins in the highly curated Uniprot Swiss-Prot database (*n*=561,176). *Ab initio* gene prediction was based on *A. umbellulata* specific and *O. sativa* hidden Markov models for Augustus and SNAP, respectively. Putative gene function was identified using BLAST searches of the predicted peptide sequences against the Swiss-Prot database using MARKER’s default cut-off values (1E– 6). Gene models were also classified as high confidence and low confidence based on predicted proteins using custom python scripts. High confidence models exhibited 1) a blastp hit (<1.0E-10) against the Magnoliopsida TrEMBL database (https://www.uniprot.org/)^59^ a query coverage and length within 25% of the subject coverage and length, and 3) > 66% percent identity between the query and subject.

#### Annotation of transposable elements and comparison of TE family proportions between U and A-B-D sub-genomes

TEs were annotated using CLARITE^60^ with the same parameters as described initially for *Triticum aestivum* Chinese Spring^30^ and other *Triticum/Aegilops*^28,61^. Briefly, TEs were predicted by similarity-search against the ClariTeRep library (which includes TREP http://botserv2.uzh.ch/kelldata/trep-db/) using RepeatMasker^57^ and were then modelled with CLARITE that resolve overlapping predictions, merge adjacent fragments of a single element, and eventually reconstruct patterns of nested insertions. Abundance of the different TE families of the U-genome was calculated by cumulating the length (in bps) of all predictions assigned to the same (sub)family by CLARITE. To investigate the specificity of the U-genome TE landscape compared to A, B, and D sub-genomes, we calculated fold-changes of proportions for each family compared to that annotated by the same approach in Chinese Spring RefSeq v2.1^30^. Abundance of each family was represented by the percentage of bps relatively to the (sub)genome size (all scaffolds for *Ae. umbellulata* genome assembly; only pseudomolecules for *T. aestivum*, i.e., not considering chrUn). Enrichment of TE families in the U-genome was investigated by computing log2 ratios between proportions observed in U *versus* A, B, or D. Only families accounting for more than 100 kb in at least one of the compared (sub)genomes was considered (i.e., 291 out of 501 families).

#### Genome, transcriptome and annotation quality assessment

Inspector v1.0.2 (Chen et al., 2021) was used to evaluate the quality of the Hi-C scaffolded assembly using the PacBio HiFi reads. Inspector efficiently identifies and can correct both small-scale miss assemblies (base correction) as well as large-scale structural assembly errors. Furthermore, the completeness of the corrected Hi-C assembly, transcriptome and annotation was assessed relative to conserved orthologous genes within Poaceae (poales_odb10) using BUSCO v5.0.0^62^ with the “long” argument which applies Augustus optimization for self-training^63^.

#### Whole genome resistance gene analog (RGA) prediction

To predict RGAs in the assembled genome, we used the docker image of RGAugury^27^ (https://bitbucket.org/yaanlpc/rgaugury/src/master/) with the default parameters and databases such as pfam and interproscan.

### Whole Genome comparison with wheat and wild wheat relatives

#### Phylogenetic analysis

We used ten species (*T. aestivum*, *T. turgidum* ssp. *dicoccoides*, *T. turgidum* ssp. *durum*, *T. urartu*, *Ae. tauschii*, *Ae. speltoides*, *Ae. longissima*, *Ae. sharonensis, Hordeum vulgare*, and *Secale cereale*) alongwith an outgroup species *B. distachyon* to perform phylogenetic analysis to find phylogenetic topologies for newly sequenced *Ae. umbellulata*. We used Orthofinder^64^ (v2.5.5) to find orthologs from high confidence genes for phylogenetic analysis. The high confidence genes from all the species were filtered using the same criteria that was used for classifying *Ae. umbellulata* gene models. The default parameters were used for Orthofinder to locate the multi-locus orthologous genes and generate species tree. The phylogenetic tree was visualized using online tree visualization tool iTol^65^ (v6.8.1) (https://itol.embl.de/).

#### Whole genome analysis and collinearity and synteny analysis

For whole genome analysis, we used minimap2^66^ (v2.24) to generate pairwise alignment between *Ae. umbelluata* and other genomes. A custom R script was used to plot the dot plots using R package ggplot2. Further, synteny analysis was performed using MCScanX^67^ (https://github.com/wyp1125/MCScanX). Blastp was used to generate the all vs all protein alignments and a custom python script was used to prepare bed files from gff files for each species. An online tool SynVisio^68^ (https://synvisio.github.io/) was used to visualize the MCScanX output and to generate synteny plots.

### Whole genome sequencing (WGS) of twenty *Ae. umbellulata* accessions

#### Plant material and Illumina short-read sequencing

We selected twenty geographically diverse *Ae. umbellulata* accessions based on the disease phenotyping scores for five wheat diseases (Supplementary table 13) for whole genome sequencing. Around 400 mg of leaf tissue was collected from a five weeks old plant in liquid nitrogen. Each accession was grown from a single seed to maintain genetic purity. The collected leaf samples were sent to Novogene for DNA extraction, library preparation, and sequencing (150 bp paired-end) at 10x coverage using the Illumina.

#### Mapping and short variant calling

After filtering out the secondary and other low quality alignments (MapQ < 20), the BAM files were sorted according to genomic coordinates and a CSI index was generated for each accession with SAMtools^69^ (v1.15). Genome Analysis Toolkit^70^ (GATK v4.1.8.1) (https://gatk.broadinstitute.org/hc/en-us) and Picard (v2.22.8) were used for SNP calling. First, index and sequence dictionary for reference *Ae. umbellulata* genome was generated using SAMtools “faidx” command and GATK CreateSequenceDictionary tool, respectively. Duplicate reads were marked with MarkDuplicatesSpark command and CSI index for each duplicate marked bam file was generated with SAMtools. These duplicated marked bam files for each accession were used for SNP calling individually using GATK HaplotypeCaller with –ERC GVCF option. GATK CombineGVCFs tool was used to combine all twenty variant called files for joint genotyping using GenotypeGVCFs tool. SNPs were pulled out from joint genotyping files using GATK SelectVariants tools. The SNPs were filtered using VariantFiltration tool with parameters (QD < 2.0, FS > 60.0, MQ < 40.00, MQRankSum < –12.5, ReadPosRankSum < –8.0, SOR > 3.0) Furthermore, SNPs clusters, where 3 SNPs appeared in 10 bp window, were also filtered. VCFtools^71^ (v0.1.14) was used to remove the filtered SNPs based on the filter tags, low and high average SNP depth (4 ≤ DP ≥ 15) and we kept only b-allelic SNPs sites, and SNPs on seven chromosomes (removed unanchored chromosomes). Further, we kept SNPs with 10% or less missingness using BCFtools^69^ (v1.15).

#### Diversity analysis and population structure

For diversity analysis, 7,184,562 filtered SNPs were used for principal component analysis (PCA) using PLINK^72^ (v1.9) with default parameters. For PCA plot, R (v4.0.3) package ggplot2 was used and color coded according to the susceptibility response to the total number of leaf rust races out of eleven tested races (susceptible to 0-3 races, susceptible to nine races, susceptible to eleven races). The individual ancestry proportions were computed with the “snmf” function of R package LEA^73^ using the entropy option. For each K value from 1 to 10 (where K is the count of ancestral populations), 10 independent runs were performed on all filtered SNPs. For phylogenetic analysis VCF file was converted to PHYLIP format using vcf2phylip (v2.8) python script (https://github.com/edgardomortiz/vcf2phylip). Further, RAxML^74^ (v8.2.12) was used to generate 500 bootstrap trees (-f a –# 500 –m GTRCAT) and maximum likelihood tree (-D). Final phylogenetic tree was constructed by merging those trees (-f b –z –t –m GTRCAT) and visualized using iTOL^65^ (v6.8.1) (https://itol.embl.de/). To compute the nucleotide diversity (pi) for all twenty accessions, group I (15 accessions), group II (3 accessions), and group III (2 accessions), we used “window-pi” option of VCFtools^71^ (v0.1.14) over 1,000,000 bp sliding window and used R package ggplot2 for plotting nucleotide diversity. Further, to check pairwise nucleotide diversity between reference genome accession and other 19 accessions, we used the same above-mentioned VCFtools parameters and for chromosome wise plotting of each comparison, python3 libraries pandas and matplotlib were used. To locate the geographical location of *Ae. umbellulata* accessions on a map, we plotted the latitude and longitude data using R package ggplot2, maps, and mapdata.

### *Lr9* exon capturing using WGS data of 20 resequenced *Ae. umbellulata* accessions

Based on the sequence similarity of cloned *Lr9* gene^35^, we filter out the reads from 20 *Ae. umbellulata* accessions mapped on the *Lr9* region in *Ae. umbellulata* assembly and visualized using JBrowse2^75^. The filtered reads from the exonic region of *Lr9* were assembled using Trinity^76^ (v2.15.1) and completeness of assembled *Lr9* alleles was checked using web based blastn. The assembled *Lr9* haplotypes were further translated to proteins using a web based tool Expasy (https://web.expasy.org/translate/). Further, to check for the amino acid substitutions in LR9 protein identified in resequenced accessions compared to cloned LR9, online blastp comparison was conducted and based on the amino acids changes, we divided the LR9 proteins in different haplotypes. Finally, we use AlphaFold^77^ to predict protein structures and we visualized and compared the protein structures using PyMol^78^.

## Data availability

The data will be available on request and deposited to NCBI server.

## Supplementary figures

**Supplementary figure 1.** (a) Hi-C contact map showing the intra-chromosomal interaction heatmap in the assembled chromosomes of *Ae. umbellulata*; **(b)** BUSCO assessments for analyzing the quality of assembled genome; **(c)** Graphical presentation of *Ae. umbellulata* chromosomes (blue bars) with telomeres (red); **(d)** Graphical presentation chloroplast and mitochondrial genomes of *Ae. umbellulata*.

**Supplementary figure 2.** Types of transposable elements (TEs) identified in *Aegilops umbellulata* in comparison to three sub-genomes of wheat (RefSeq_v2.1): **(a)** total, class-I, class-II, and unclassified TEs; **(b)** different types of class-I TEs; and **(c)** different types of class-II TEs.

**Supplementary figure 3.** Comparative genome analysis of *Aegilops umbellulata* with *Brachypodium distachyon* **(a)**, *Hordeum vulgare* (H) **(b)**, and A, B, and D sub-genomes of *Triticum aestivum* **(c)**.

**Supplementary figure 4.** Syntenic analysis of *Aegilops umbellulata* (au) with *Triticum urartu* (tu), *Aegilops speltoides* (as), and *Aegilops sharonensis* (sh).

**Supplementary figure 5.** Syntenic analysis of individual *Aegilops umbellulata* (au) chromosomes with *Aegilops tauschii* (at) and *Aegilops longissima* (al).

**Supplementary figure 6.** Disease reaction of Aegilops umbellulata accessions and susceptible wheat cultivars (Prosper, Thatcher, and Morocco) to United States most virulent races, TNBJS **(a)** and MNPSD **(b)** of leaf rust.

**Supplementary figure 7.** Bacterial leaf streak (BLS) resistance in *Aegilops umbellulata* accession, PI 554417 compared to sequenced accession, PI 554389.

**Supplementary figure 8.** Morphological variation for spike architecture in *Aegilops umbellulata*.

**Supplementary figure 9.** Morphological variation for plant architecture and flowering in *Aegilops umbellulata*.

**Supplementary figure 10.** Nucleotide diversity (π) analysis of 20 *Aegilops umbellulata* accessions.

**Supplementary figure 11.** Multiple sequence alignment of cloned *Lr9* gene from TA1851 and six *Lr9* haplotypes (Hap) found in 20 *Aegilops umbellulata* accessions.

## Supplementary tables

**Supplementary table 1.** Raw data yield of PacBio HiFi CCS sequencing.

**Supplementary table 2.**Raw data yield of Nanopore MinION sequencing.

**Supplementary table 3.** Summary statistics of genome assembly using PacBio HiFi reads.

**Supplementary table 4.** Summary statistics of scaffolds after chromosome conformation capture (Hi-C).

**Supplementary table 5.** Information about manual gap closing.

**Supplementary table 6.** Synteny guided chromosome assignment to scaffolds based on IWGSC RefSeq_v2.1 D sub-genome.

**Supplementary table 7.** Summary of chromosome length, genomic content, and predicted gene models of *Aegilops umbellulata* genome assembly.

**Supplementary table 8.** Type and number of predicted resistance gene analogs (RGAs) in *Aegilops umbellulata*.

**Supplementary table 9.** Comparison of GC content (%) of *Aegilops umbellulata* and wheat reference genome.

**Supplementary table 10.** Genome completeness of *Aegilops umbellulata* based on BUSCO score.

**Supplementary table 11.** Proportion of Transposable Elements (TE) at whole genome level for *Aegilops umbellulata* and wheat reference (v2.1) genome.

**Supplementary table 12.** High and Low confidence gene models for different species used in phylogenetic analysis.

**Supplementary table 13.** Disease phenotyping data for leaf rust, stripe rust, stem rust, bacterial leaf streak (BLS), and tan spot on *Aegilops umbellulata* diverse panel.

**Supplementary table 14.** Information about the number of retained single nucleotide polymorphisms (SNPs) after each filtering step.

**Supplementary table 15.** Information about amino acid changes in *Lr9* alleles/haplotypes found in *Aegilops umbellulata* compared to cloned *Lr9*.

**Supplementary table 16.** Positions of amino acid changes in *Lr9* gene in *Aegilops umbellulata* accessions compared to cloned *Lr9*.

## Declarations

The authors declare no competing interests.

## Acknowledgments

This research was funded in part by USDA National Institute of Food and Agriculture (NIFA; Project no. 2023-67014-39347), USDA-ARS CRIS project number 3060-2100-046-000D, and the U.S. Department of Agriculture–National Institute of Food and Agriculture (Hatch Project ND02243). Authors acknowledge the support provided by the Agricultural Experiment Station and Department of Plant Pathology at North Dakota State University, Fargo, ND, USA.

