## Supplementary Figures for "Genomes of *Aegilops umbellulata* provide new insights into unique structural variations and genetic diversity in the U-genome for wheat improvement"

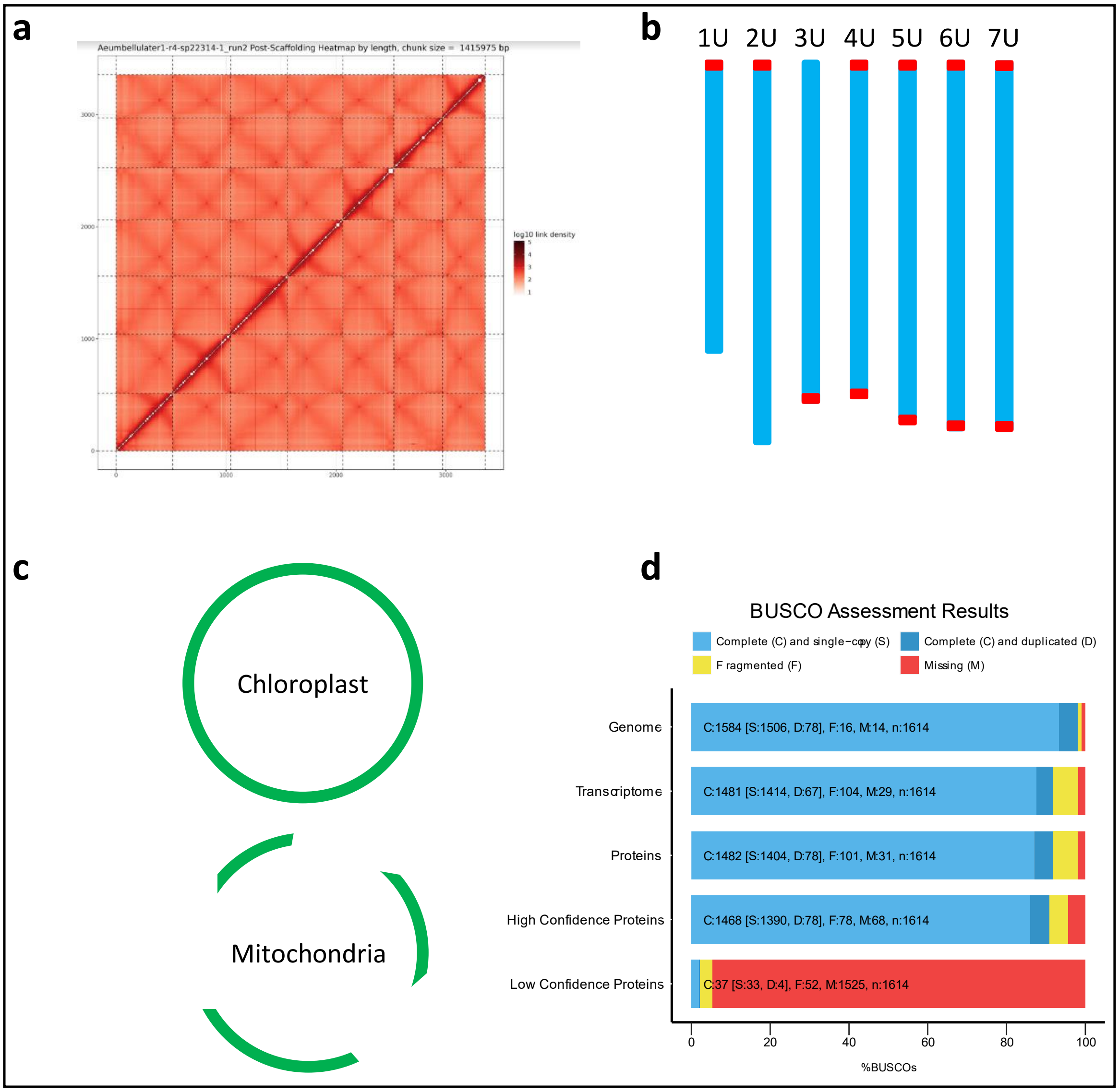


**Supplementary figure 1**: **(a)** Hi-C contact map showing the intrachromosomal interaction heatmap in the assembled chromosomes of *Ae. umbellulata*; **(b)** Graphical presentation of *Ae. umbellulata* chromosomes (blue bars) with telomeres (red); **(c)** Graphical presentation chloroplast and mitochondrial genomes of *Ae. umbellulata*; **(d)** BUSCO assessments for analyzing the quality of assembled genome.

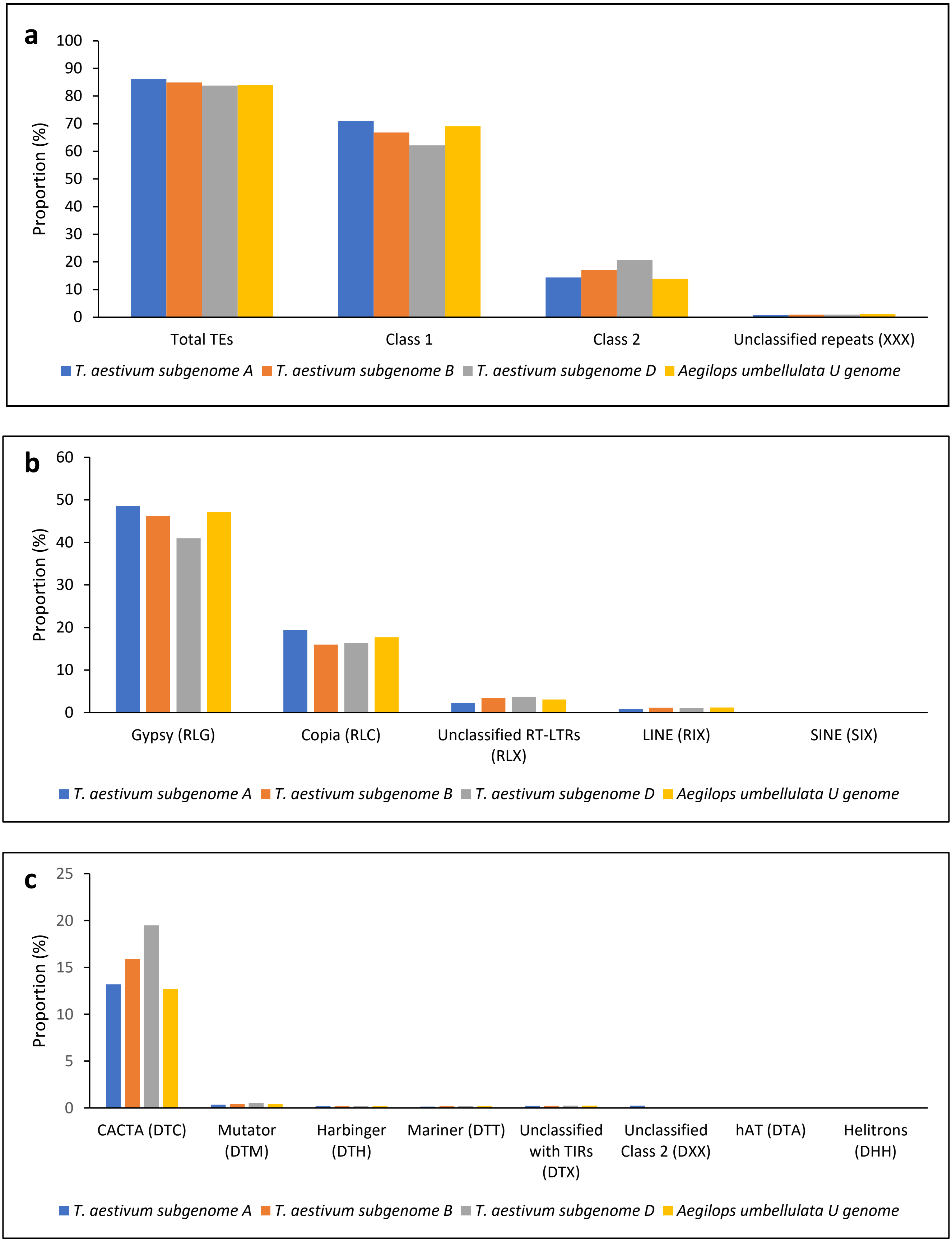


**Supplementary figure 2**: Types of transposable elements (TEs) identified in *Aegilops umbellulata* in comparison to three sub-genomes of wheat (ref v2.1). **(a)** Proportion of class-I, class-II, and unclassified TEs; **(b)** proportion of sub-classes within class-I TEs; and **(c)** proportion of sub-classes of class-II TEs.

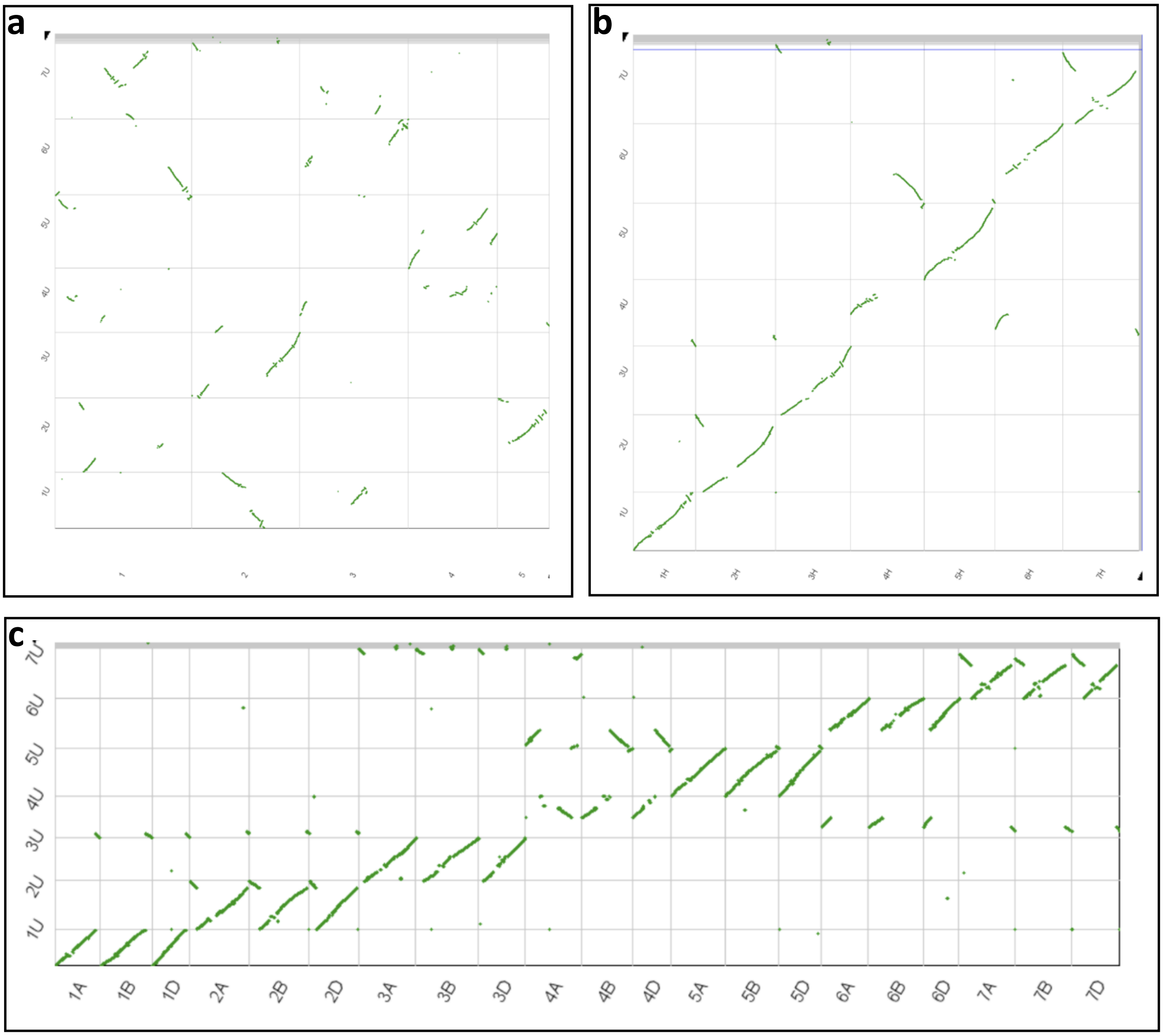


**Supplementary figure 3**: Comparative genome analysis of *Aegilops umbellulata* with *Brachypodium distachyon* **(a)**, *Hordeum vulgare* **(b),** and A-B-D sub-genomes of *Triticum aestivum* **(c).**

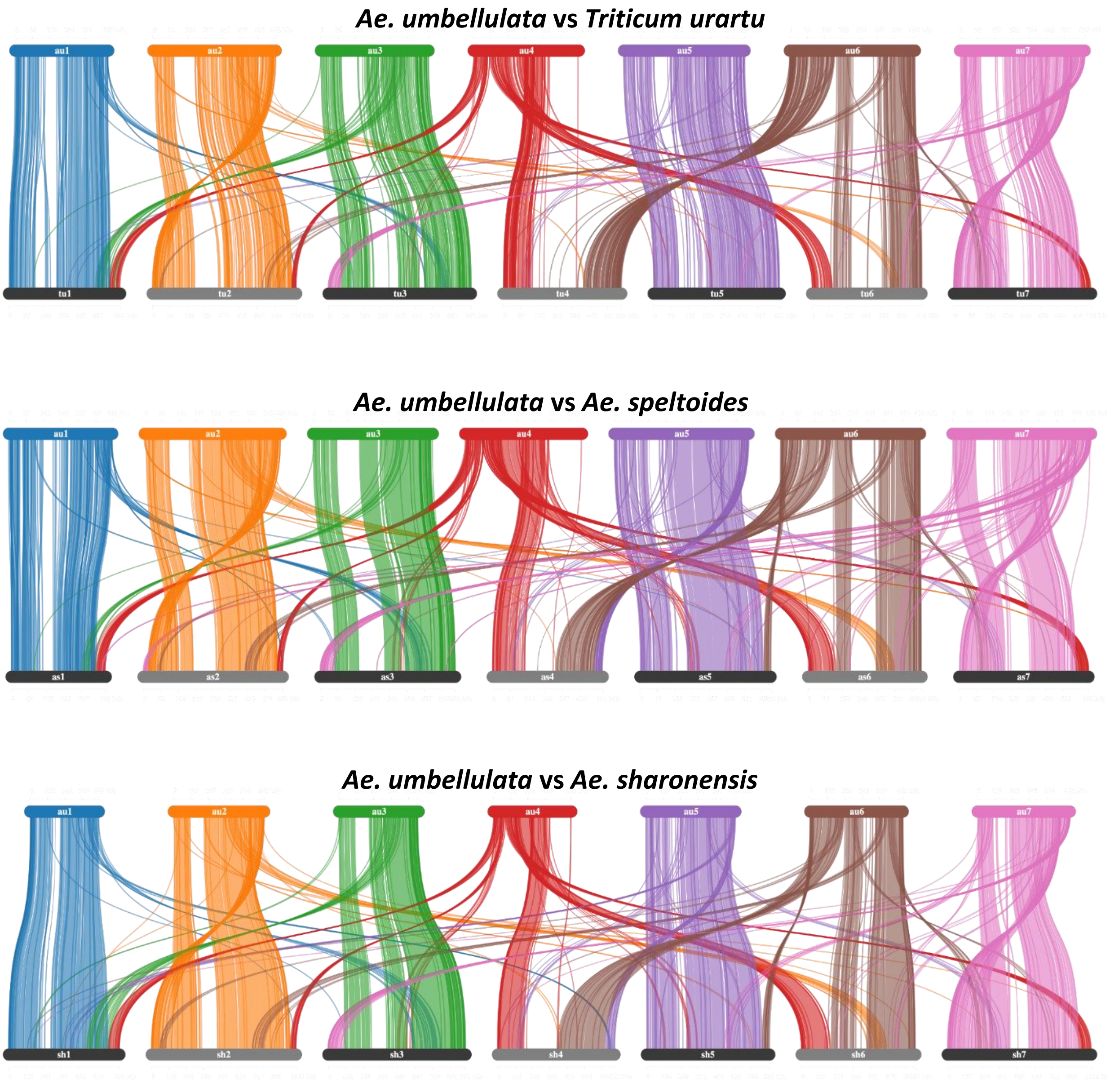


**Supplementary figure 4**: Syntenic relationship of *Aegilops umbellulata* (au) with *Triticum urartu* (tu), *Aegilops speltoides* (as), and *Aegilops sharonensis* (sh).

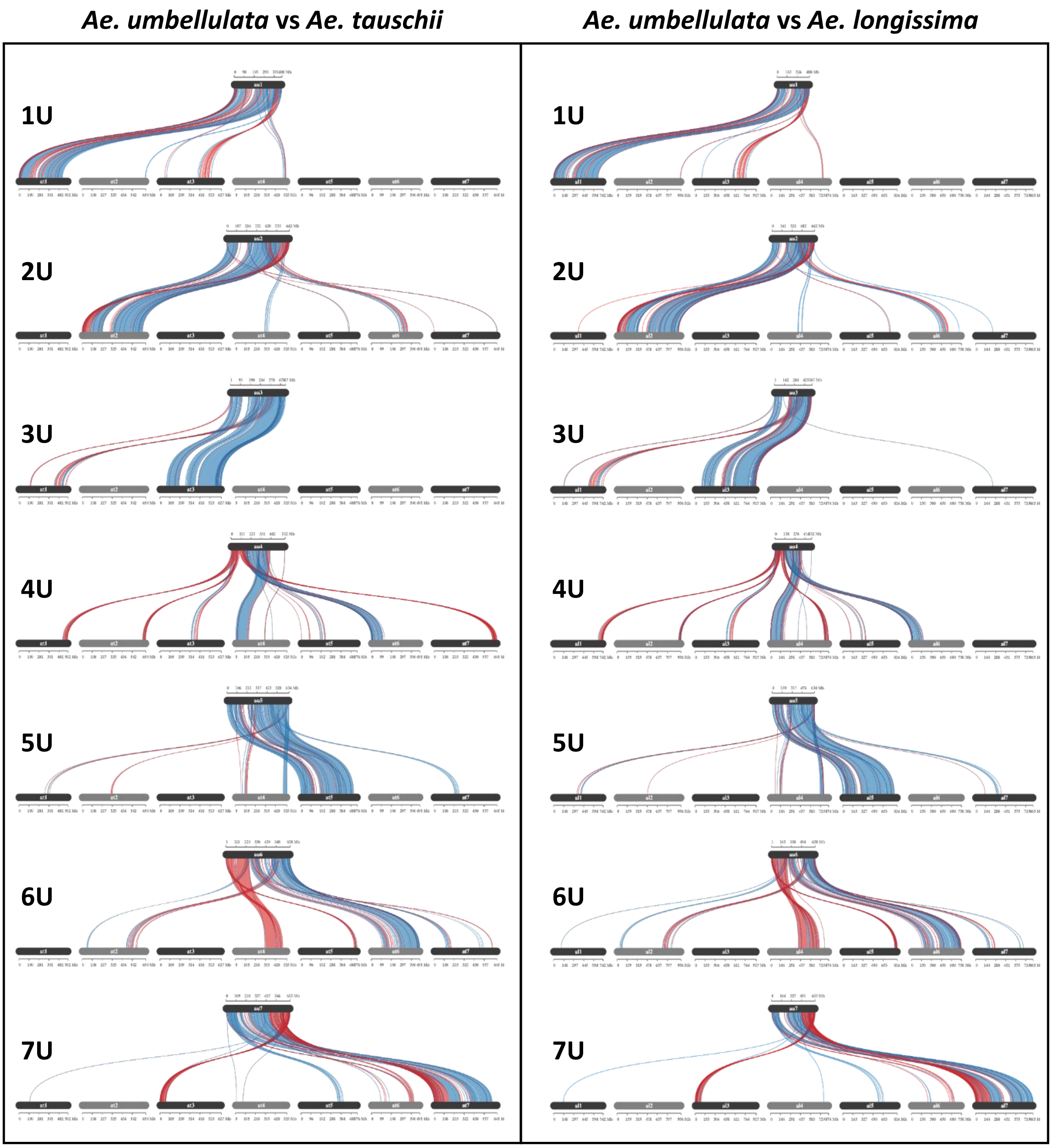


**Supplementary figure 5**: Syntenic relationship of individual *Aegilops umbellulata* (au) chromosomes with *Aegilops tauschii* (at) and *Aegilops longissima* (al).

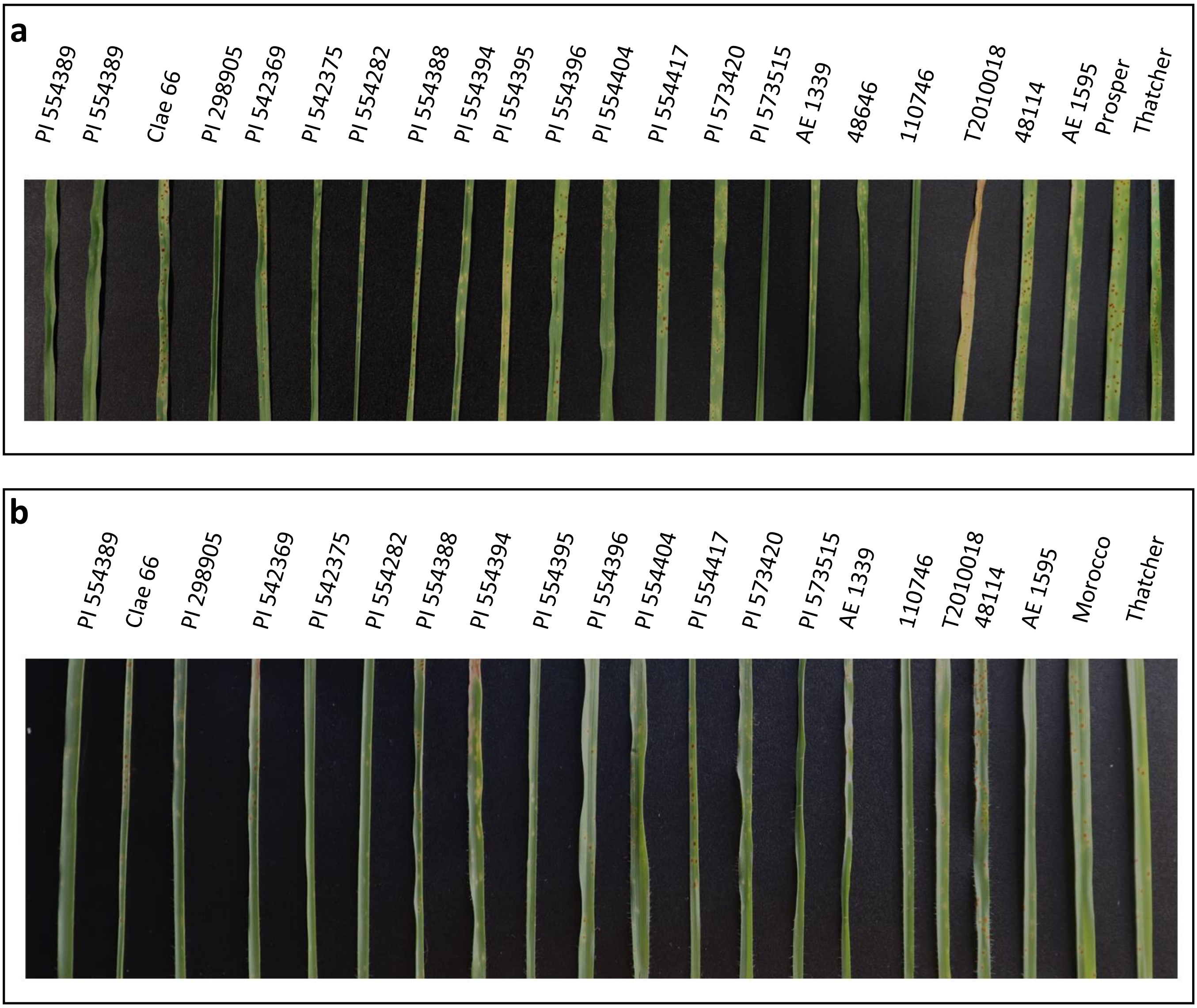


**Supplementary figure 6**: Disease reaction of *Aegilops umbellulata* accessions and susceptible wheat cultivars (Prosper, Thatcher, and Morocco) to the United States *Puccinia triticina* races, **(a)** TNBJS and **(b)** MNPSD.

**PI 554417** **PI 554389**

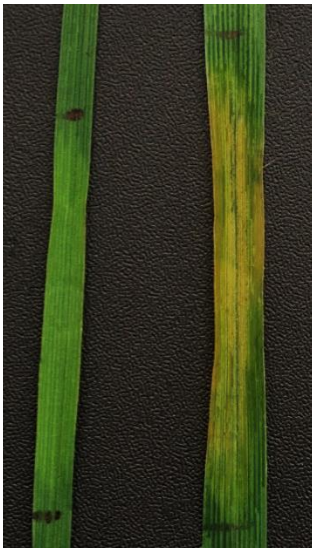

**Supplementary figure 7**: Bacterial leaf streak (BLS) resistance in *Aegilops umbellulata* accession, PI 554417 compared to sequenced accession, PI 554389 against BLS-P3 isolate at seedling stage.

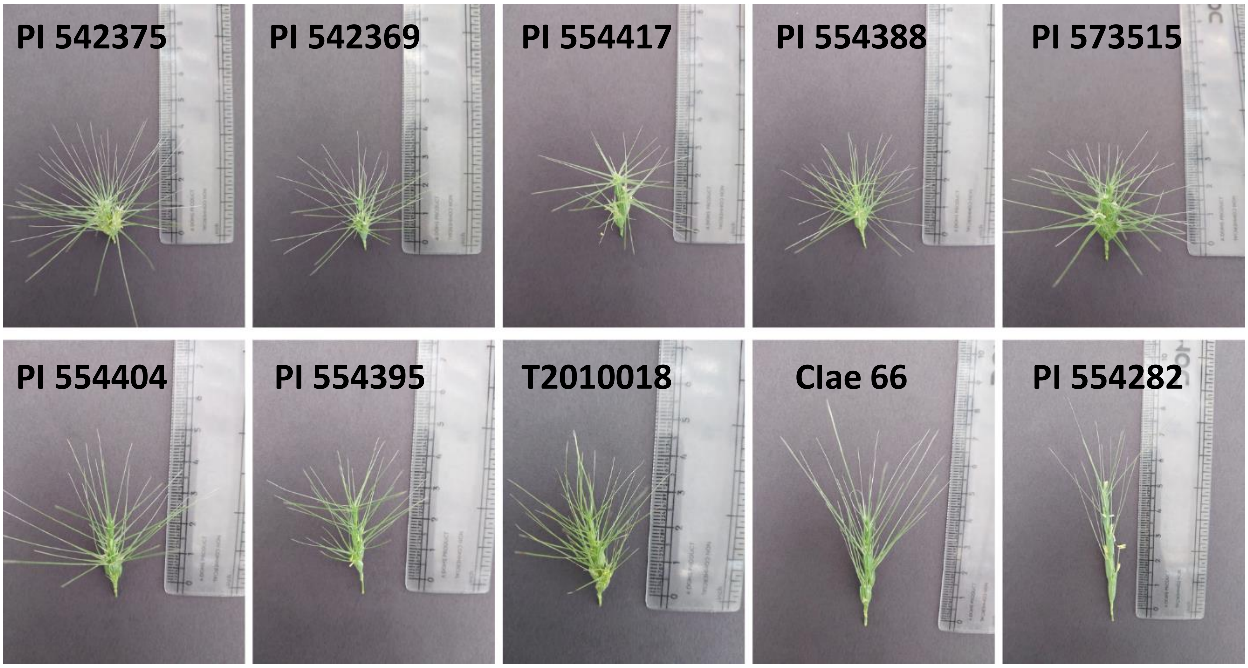


**Supplementary figure 8**: Morphological variation for spike architecture in *Aegilops umbellulata.*

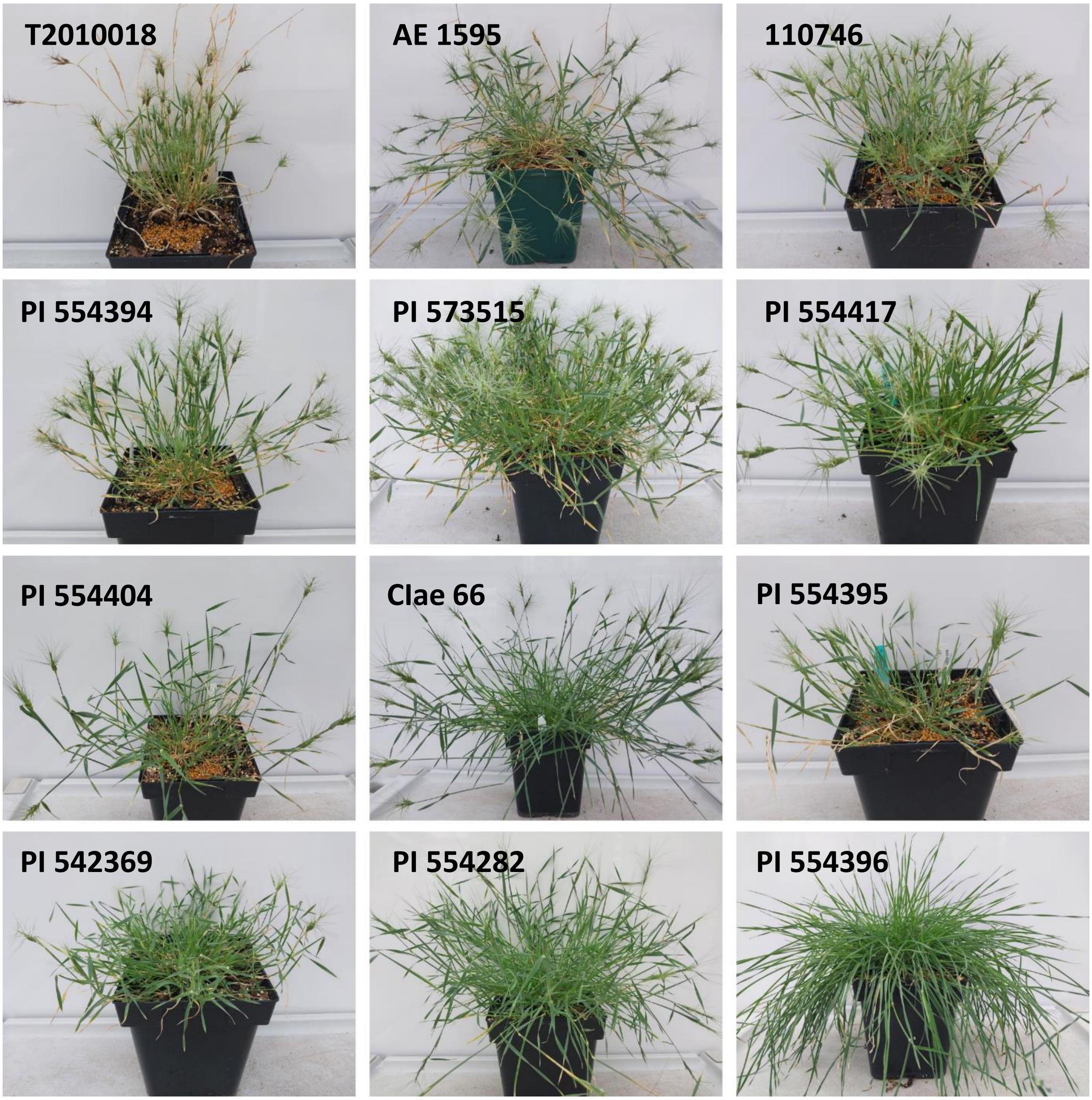


**Supplementary figure 9**: Morphological variation for plant architecture and flowering in *Aegilops umbellulata.*

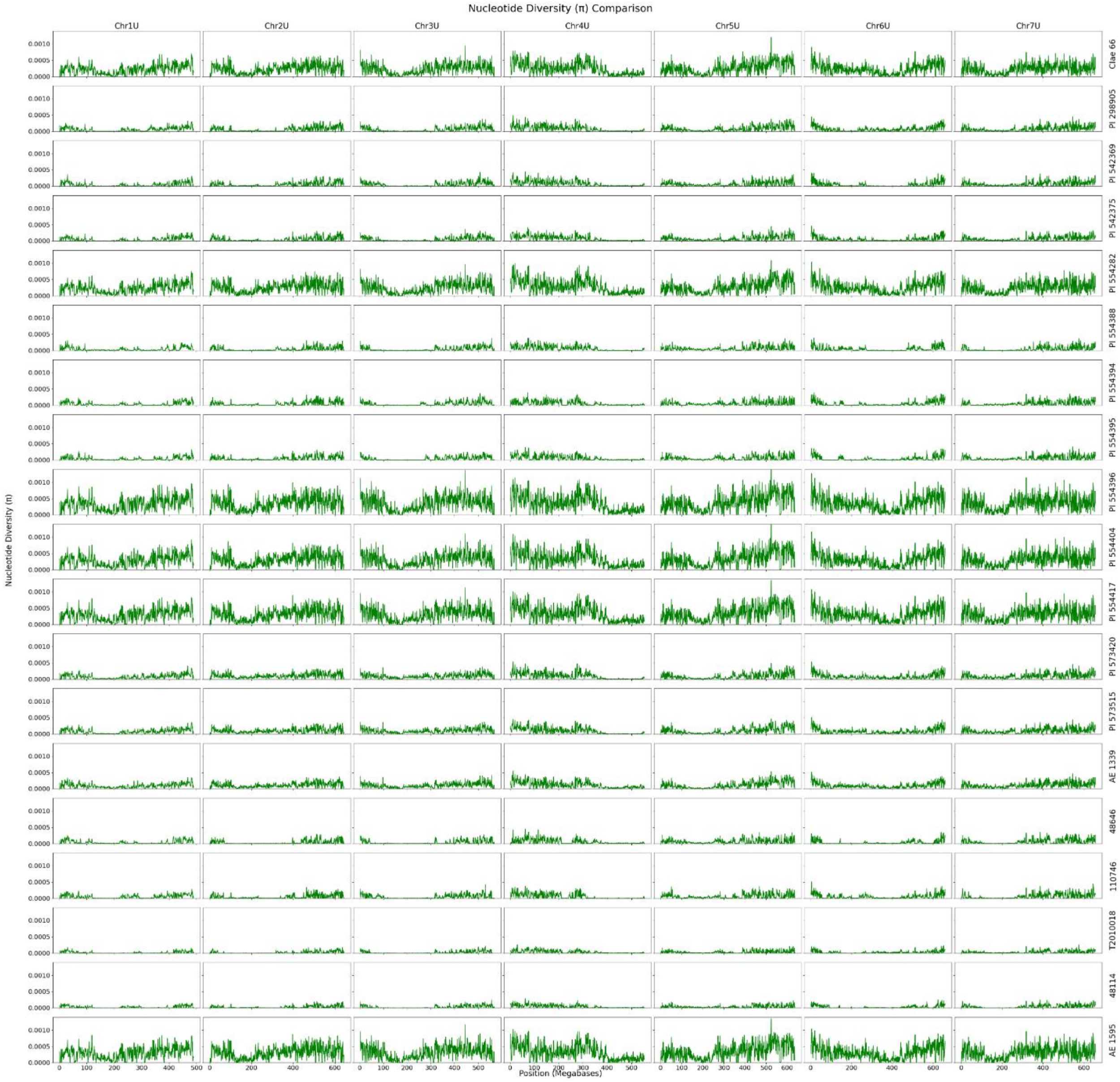


**Supplementary figure 10**: Pairwise nucleotide diversity ($\pi$) analysis of resequenced *Aegilops umbellulata* accessions compared to sequenced PI 554389 for each chromosome.

```
          1        10        20         30        40        50
1 Lr9     MASPHGSDGAPKSEDLQLSSS.SSNSSTNLSSSSSSSSSSLSSRSSIDSTMGSKLQKAK
2 Hap_1   MASPHGSDGAPKSEDLQLSSS.SSNSSTNLSSSSSSSSSSLSSRSSIDSTMGSKLQKAK
3 Hap_2   MASPHGSDGAPKSEDLQLSSSTSSNSSTNLSSSSSSSSSSLSSRSSIDSTMGSKLQKAK
4 Hap_3   MASPHGSDGAPKSEDLQLSSS.SSNSSTNLSSSSSSSSSSLSSRSSIDSTMGSKLQKAK
5 Hap_4   MASPHGSDGAPKSEDLQLSSS.SSNSSTNLSSSSSSSSSSLSSRSSIDSTMGSKLQKAK
6 Hap_5   MASPHGSDGAPKSEDLQLSSS.SSNSSTNLSSSSSSSSSSLSSRSSIDSTMGSKLQKAK
7 Hap_6   MASPHGSDGAPKSEDLQLSSS.SSNSSTNLSSSSSSSSSSLSSRSSIDSTMGSKLQKAK
```

```
         60        70        80        90        100       110
1 Lr9     RLSIEHLEVITDNFSQERVIGRGGFGIVYKGVHANGKNIAVKVLHKNMTGTDDGQFEHEF
2 Hap_1   RLSIEHLEVITDNFSQERVIGRGGFGIVYKGVHANGKNIAVKVLHKNMTGTDDGQFEHEF
3 Hap_2   RLSIEHLEVITDNFSQERVIGRGGFGIVYKGVHANGKNIAVKVLHKNMTGTDDGQFEHEF
4 Hap_3   RLSIEHLEVITDNFSQERVIGRGGFGIVYKGVHANGKNIAVKVLHKNMTGTDDGQFEHEF
5 Hap_4   RLSIEHLEVITDNFSQERVIGRGGFGIVYKGVHANGKNIAVKVLHKNMTGTDDGQFEHEF
6 Hap_5   RLSIEHLEVITDNFSQERVIGRGGFGIVYKGVHANGKNIAVKVLHKNMTGTDDGQFEHEF
7 Hap_6   RLSIEHLEVITDNFSQERVIGRGGFGIVYKGVHANGKNIAVKVLHKNMTGTDDGQFEHEF
```

```
        120       130       140       150       160       170
1 Lr9     KNLMRLKHQNIVQLVGYCYETQHKPMLHEGVAIFADDIYKALCFEYMENRSLSKHISVEC
2 Hap_1   KNLMRLKHQNIVQLVGYCYETQHKPMLHEGVAIFADDIYKALCFEYMENRSLSKHISVEC
3 Hap_2   KNLMRLKHQNIVQLVGYCYETQHKPMLHEGVAIFADDIYKALCFEYMENRSLSKHISVEC
4 Hap_3   KNLMRLKHQNIVQLVGYCYETQHKPMLHEGVAIFADDIYKALCFEYMENRSLSKHISVEC
5 Hap_4   KNLMRLKHQNIVQLVGYCYETQHKPMLHEGVAIFADDIYKALCFEYMENRSLSKHISVEC
6 Hap_5   KNLMRLKHQNIVQLVGYCYETQHKPMLHEGVAIFADDIYKALCFEYMENRSLSKHISVEC
7 Hap_6   KNLMRLKHQNIVQLVGYCYETQHKPMLHEGVAIFADDIYKALCFEYMENRSLSKHISVEC
```

```
        180       190       200       210       220       230
1 Lr9     DGLEWHTRYKIIKGTCEGLKYLHEEFKEPIYHLDLKPDNILLDENMVPKLADFGSSKLFG
2 Hap_1   DGLEWHTRYKIIKGTCEGLKYLHEEFKEPIYHLDLKPDNILLDENMVPKLADFGSSKLFG
3 Hap_2   DGLEWHTRYKIIKGTCEGLKYLHEEFKEPIYHLDLKPDNILLDENMVPKLADFGSSKLFG
4 Hap_3   DGLEWHTRYKIIKGTCEGLKYLHEEFKEPIYHLDLKPDNILLDENMVPKLADFGSSKLFG
5 Hap_4   DGLEWHTRYKIIKGTCEGLKYLHEEFKEPIYHLDLKPDNILLDENMVPKLADFGSSKLFG
6 Hap_5   DGLEWHTRYKIIKGTCEGLKYLHEEFKEPIYHLDLKPDNILLDENMVPKLADFGSSKLFG
7 Hap_6   DGLEWHTRYKIIKGTCEGLKYLHEEFKEPIYHLDLKPDNILLDENMVPKLADFGSSKLFG
```

```
        240       250       260       270       280       290
1 Lr9     KELTMVITQNFTATIGYMPPEFISHQIVSKKFDIFSLGVVMTKIIAGPRGNSLSFEMDYQ
2 Hap_1   KELTMVITQNFTATIGYMPPEFISHQIVSKKFDIFSLGVVMTKIIAGPRGNSLSFEMDYQ
3 Hap_2   KELTMVITQNFTATIGYMPPEFISHQIVSKKFDIFSLGVVMTKIIAGPRGNSLSFEMDYQ
4 Hap_3   KELTMVITQNFTATIGYMPPEFISHQIVSKKFDIFSLGVVMTKIIAGPRGNSLSFEMDYQ
5 Hap_4   KELTMVITQNFTATIGYMPPEFISHQIVSKKFDIFSLGVVMTKIIAGPRGNSLSFEMDYQ
6 Hap_5   KELTMVITQNFTATIGYMPPEFISHQIVSKKFDIFSLGVVMTKIIAGPRGNSLSFEMDYQ
7 Hap_6   KELTMVITQNFTATIGYMPPEFISHQIVSKKFDIFSLGVVMTKIIAGPRGNSLSFEMDYQ
```

```
        300       310       320       330       340       350
1 Lr9     DFLDQVQKNWRNRLQDTWHNSEPVQAYCRQVNVCIEIALSCMDADRNKRPSILDIIDKLN
2 Hap_1   DFLDQVQKNWRNRLQDTWHNSEPVQAYCRQVNVCIEIALSCMDADRNKRPSILDIIDKLN
3 Hap_2   DFLDQVQKNWRNRLQDTWHNSEPVQAYCRQVNVCIEIALSCMDADRNKRPSILDIIDKLN
4 Hap_3   DFLDQVQKNWRNRLQDTWHNSEPVQAYCRQVNVCIEIALSCMDADRNKRPSILDIIDKLN
5 Hap_4   DFLDQVQKNWRNRLQDTWHNSEPVQAYCRQVNVCIEIALSCMDADRNKRPSILDIIDKLN
6 Hap_5   DFLDQVQKNWRNRLQDTWHNSEPVQAYCRQVNVCIEIALSCMDADRNKRPSILDIIDKLN
7 Hap_6   DFLDQVQKNWRNRLQDTWHNSEPVQAYCRQVNVCIEIALSCMDADRNKRPSILDIIDKLN
```

```
        360       370       380       390       400       410
1 Lr9     KTETVIEQLHRDHQWTSRRITMGDDKLKNEATPLKEKKSEKATEPTSLTIQVLKDITNNF
2 Hap_1   KTETVIEQLHRDHQWTSRRITMGDDKLKNEATPLKEKKSEKATEPTSLTIQVLKDITNNF
3 Hap_2   KTETVIEQLHRDHQWTSRRITMGDDKLKNEATPLKEKKSEKATEPTSLTIQVLKDITNNF
4 Hap_3   KTETVIEQLHRDHQWTSRRITMGDDKLKNEATPLKEKKSEKATKPTSLTIQVLKDITNNF
5 Hap_4   KTETVIEQLHRDHQWTSRRITMGDDKLKNEATPLKEKKSEKATKPTSLTIQVLKDITNNF
6 Hap_5   KTETVIEQLHRDHQWTSRRITMGDDKLKNEATPLKEKKSEKATEPTSLTIQVLKDITNNF
7 Hap_6   KTETVIEQLHRDHQWTSRRITMGDDKLKNEATPLKEKKSEKATEPTSLTIQVLKDITNNF
```

```
          420       430       440       450       460       470
1 Lr9     AEERVIGRGAYGTVYMGKHENGEEIAVKLLYNNMQLVDDEQFENDFYGLMMLNHPNIVRL
2 Hap_1   AEERVIGRGAYGTVYMGKHENGEEIAVKLLYNNMQLVDDEQFENDFYGLMMLNHPNIVRL
3 Hap_2   AEERVIGRGAYGTVYMGKHENGEEIAVKLLYNNMQLVDDEQFENDFYGLMMLNHPNIVRL
4 Hap_3   AEERLIGRGAYGRVYMGKHENGEKIAVKLLNNNMQVVDDEQFKYEFYGLMRFNHPNIVRL
5 Hap_4   AEERLIGRGAYGRVYMGKHENGEKIAVKLLNNNMQVVDDEQFKYEFYGLMRFNHPNIVRL
6 Hap_5   AEERVIGRGAYGTVYMGKHENGEEIAVKLLYNNMQLVDDEQFENDFYGLMMLNHPNIVRL
7 Hap_6   AQERVIGQGAYGKVYMGKHENGEEIAVKLLHNNMQLVDDEQFQHELCSLMMLNHPNIVRL

          480       490       500       510       520       530
1 Lr9     VGYCYETQHLPVDFQGRTVLAQTTHRALCLEYMHMGSLQRHLSDESSGPDWPTRYKIIKG
2 Hap_1   VGYCYETQHLPVDFQGRTVLAQTTHRALCLEYMHMGSLQRHLSDESSGPDWPTRYKIIKG
3 Hap_2   VGYCYETQHLPVDFQGRTVLAQTTHRALCLEYMHMGSLQRHLSDESSGPDWPTRYKIIKG
4 Hap_3   VGYCYETQRQHTIFQGRIVFAETTYRALCLEYMHMGSLQRHLSDESSGPDWPTRYKIIKG
5 Hap_4   VGYCYETQRQHTIFQGRIVFAETTYRALCLEYMHMGSLQRHLSDESSGPDWPTRYKIIKG
6 Hap_5   VGYCYETQHLPVDFQGRTVLAQTTHRALCLEYMHMGSLQRHLSDESSGPDWPTRYKIIKG
7 Hap_6   VGYCYETQHLPVDFQGRTVLAQKTHRALCLEYMHMGSLQRHLSDESSGPDWPTRYKIIKG

          540       550       560       570       580       590
1 Lr9     TCEGLKYLHEGLDKPLYHLDLKPENILLDKNMMPKLADFGLTKLFGEEQTRVTHSLFGTM
2 Hap_1   TCEGLKYLHEGLDKPLYHLDLKPENILLDKNMMPKLADFGLTKLFGEEQTRVTHSLFGTM
3 Hap_2   TCEGLKYLHEGLDKPLYHLDLKPENILLDKNMMPKLADFGLTKLFGEEQTRVTHSLFGTM
4 Hap_3   TCEGLKYLHEGLDKPLYHLDLKPENILLDKNMMPKLADFGLTKLFGEEQTRVTHSLFGTM
5 Hap_4   TCEGLKYLHEGLDKPLYHLDLKPENILLDKNMMPKLADFGLTKLFGEEQTRVTHSLFGTM
6 Hap_5   TCEGLKYLHEGLDKPLYHLDLKPENILLDKNMMPKLADFGLTKLFGEEQTRVTHSLFGTM
7 Hap_6   TCEGLKYLHEGLDKPLYHLDLKPENILLDKNMMPKLADFGLTKLFGEEQTRVTHSLFGTM

          600       610       620       630       640       650
1 Lr9     GYLPPEFISHQVVSKKFDIFSLGVVMTKIIAGPMGYHMSVEMLEEDFVDQVHEKWRNRME
2 Hap_1   GYLPPEFISHQVVSKKFDIFSLGVVMTKIIAGPMGYHMSVEMPEEDFVDQVHEKWRNRME
3 Hap_2   GYLPPEFISHQVVSKKFDIFSLGVVMTKIIAGPMGYHMSVEMPEEDFVDQVHEKWRNRME
4 Hap_3   GYLPPEFISHQVVSKKFDIFSLGVVMTKIIAGPMGYHMSVEMPEEDFVDQVHEKWRNRME
5 Hap_4   GYMPPEFISHQVVSKKFDIFSLGVVMTKIIAGPMGYHMSVEMPEEDFVDQVHEKWRNRME
6 Hap_5   GYLPPEFISHQVVSKKFDIFSLGVVMTKIIAGPMGYHMSVEMPEEDFVDQVHEKWRNRME
7 Hap_6   GYLPPEFISHQVVSKKFDIFSLGVVMTKIIAGPMGYHMSAEMPEEDFVDQVHEKWRNRLE

          660       670       680       690       700       710
1 Lr9     ATCTSSQTLDAYCEQVRICTKIGLSCVEYDRHKRPNILDIIDRLDETERKIEKALFSLSD
2 Hap_1   ATCTSSQTLDAYCEQVRICTKIGLSCVEYDRHKRPNILDIIDRLDETERKIEKALFSLSD
3 Hap_2   ATCTSSQTLDAYCEQVRICTKIGLSCVEYDRHKRPNILDIIDRLDETERKIEKALFSLSD
4 Hap_3   ATCTSSQTLDAYCEQVRICTKIGLSCVEYDRHKRPNILDIIDRLDETERKIEKALFSLSD
5 Hap_4   ATCTSSQTLDAYCEQVRICTKIGLSCVEYDRHKRPNILDIIDRLDETERKIEKALFSLSD
6 Hap_5   ATCTSSQTLDAYCEQVRICTKIGLSCVEYDRHKRPNILDIIDRLDETERKIEKALFSLSD
7 Hap_6   ATCTSSQTLDAYCEQVRICTKIGLSCVEYDRHKRPNILDIIDRLDETERKIEKALFSLSD

          720       730       740       750       760       770
1 Lr9     SMHSWLRGDLLKLEAFTKSQEIPSTETCNEFPVLLRITGTPWCGLEGMPRAGVDVVVVVE
2 Hap_1   SMHSWLRGDLLKLEAFTKSQEIPSTETCNEFPVLLRITGTPWCGLEGMPRAGVDVVVVVE
3 Hap_2   SMHSWLRGDLLKLEAFTKSQEIPSTETCNEFPVLLRITGTPWCGLEGMPRAGVDVVVVVE
4 Hap_3   SMHSWLRGDLLKLEAFTKSQEIPSTETCNEFPVLLRITGTPWCGLEGMPRAGVDVVVVVE
5 Hap_4   SMHSWLRGDLLKLEAFTKSQEIPSTETCNEFPVLLRITGTPWCGLEGMPRAGVDVVVVVE
6 Hap_5   SMHSWLRGDLLKLEAFTKSQEIPSTETCNEFPVLLRITGTPWCGLEGMPRAGVDVVVVVE
7 Hap_6   SMHSWLRGDLLKLEAFTKSQEIPSTETCNEFPVLLRITGTPWCGLEGMPRAGVDVVVVVE

          780       790       800       810       820       830
1 Lr9     VDWFMVLQWRLDIIKQALTVVIDKLGPTDRLSILSFEGDVPHIMKLAFMSDQGRDAAKLV
2 Hap_1   VDWFMVLQWRLDIIKQALTVVIDKLGPTDRLSILSFEGDVPHIMKLAFMSDQGRDAAKLV
3 Hap_2   VDWFMVLQWRLDIIKQALTVVIDKLGPTDRLSILSFEGDVPHIMKLAFMSDQGRDAAKLV
4 Hap_3   VDWFMVLQWRLDIIKQALTVVIDKLGPTDRLSILSFEGDVPHIMKLAFMSDQGRDAAKLV
5 Hap_4   VDWFMVLQWRLDIIKQALTVVIDKLGPTDRLSILSFEGDVPHIMKLAFMSDQGRDAAKLV
6 Hap_5   VDWFMVLQWRLDIIKQALTVVIDKLGPTDRLSILSFEGDVPHIMKLAFMSDQGRDAAKLV
7 Hap_6   VDWFMVLQWRLDIIKQALTVVIDKLGPTDRLSILSFEGDVPHIMKLAFMSDQGRDAAKLV
```

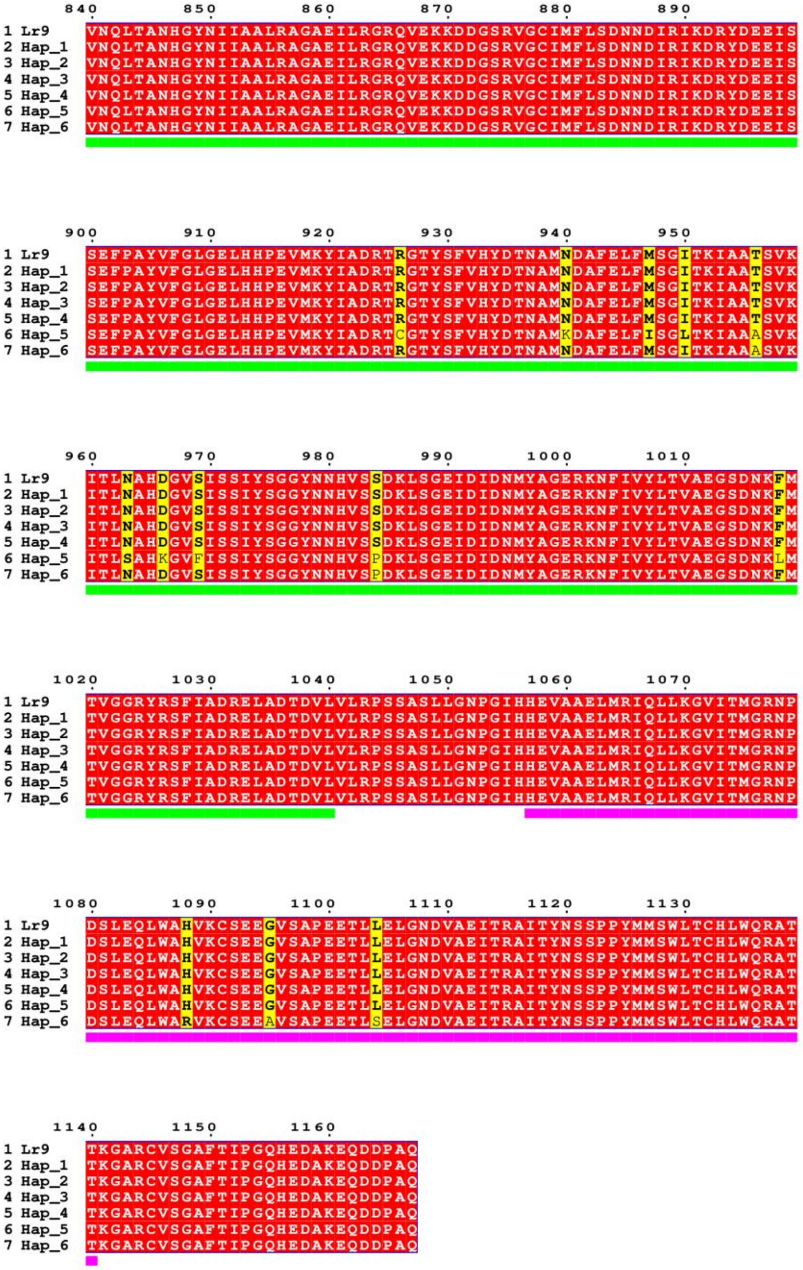


**Supplementary figure 11**: Multiple sequence alignment of cloned *Lr9* gene from TA1851 and six *Lr9* haplotypes (Hap) found in 20 *Aegilops umbellulata* accessions.
